# THE DUAL-SPECIFICITY KINASE DYRK1A INTERACTS WITH THE HEPATITIS B VIRUS GENOME AND REGULATES THE PRODUCTION OF VIRAL RNA

**DOI:** 10.1101/2024.05.30.596758

**Authors:** Florentin Pastor, Emilie Charles, Chiara Di Vona, Maëlys Chapelle, Michel Rivoire, Guillaume Passot, Benoit Chabot, Susana de la Luna, Julie Lucifora, David Durantel, Anna Salvetti

## Abstract

The genome of Hepatitis B virus (HBV) persists in infected hepatocytes as a nuclear episome (cccDNA) that is responsible for the transcription of viral genes and viral rebound, following antiviral treatment arrest in chronically infected patients. Beside pegylated interferon-alpha, there is currently no clinically approved therapeutic protocol able to target cccDNA [1]. The development of alternative strategies aiming at permanently abrogating HBV RNA production requires a thorough understanding of cccDNA transcriptional and post-transcriptional regulation. In a previous study, we discovered that 1C8, a compound that inhibits the phosphorylation of some cellular RNA-binding proteins (RBPs), could decrease the level of HBV RNAs. Here, we aimed at identifying kinases responsible for this effect. Among the kinases targeted by 1C8, we focused on DYRK1A, a dual-specificity kinase that controls the transcription of cellular genes by phosphorylating transcription factors, histones, chromatin regulators as well as RNA polymerase II. The results of a combination of genetic and chemical approaches using HBV-infected hepatocytes, indicated that DYRK1A positively regulates the production of HBV RNAs. In addition, we found that DYRK1A associates with cccDNA, and stimulates the production of HBV nascent RNAs. Finally, reporter gene assays showed that DYRK1A up-regulates the activity of the HBV enhancer 1/X promoter in a sequence-dependent manner. Altogether, these results indicate that DYRK1A is a proviral factor that may participate in the the HBV life cycle by stimulating the production of HBx, a viral factor absolutely required to trigger the complete cccDNA transcriptional program.

## INTRODUCTION

Chronic infection by Hepatitis B virus (HBV) is due to the stable persistence and activity of the viral genome, called cccDNA (for covalently-closed circular DNA), within the nucleus of infected hepatocytes. Clinically accepted treatments are represented mostly by nucleoside analogs, which target viral reverse-transcription and reduce viremia, but rarely lead to a stable functional cure. Indeed, these compounds do not affect cccDNA stability and transcriptional activity. As a consequence, the continuous production of viral RNAs and proteins allows viral replication to resume off-treatment, and, more importantly, contributes to immune escape mechanisms and liver disease progression [2,3]. Despite some experimental proof-of-concept approaches [1,4], *in vivo* therapies aiming at eradicating HBV cccDNA and other replication intermediates from infected hepatocytes are unrealistic to date [1]. Therefore, strategies directed to permanently silence the viral genome and/or destabilize the viral RNAs, requiring an in-depth understanding of cccDNA transcriptional and post-transcriptional regulation, are currently investigated.

After binding to its main receptor and co-receptors on the hepatocyte membrane, HBV particles are internalized within endosomes from which, the viral nucleocapsids are released later on. Subsequently, these nucleocapsids are directed to the nuclear membrane, translocate through the nuclear pore, and finally disassemble within the inner nuclear basket [5,6]. This process results in the release into the nucleoplasm of the viral relaxed circular DNA genome (rcDNA), with a covalently-attached polymerase, and of 240 copies of the HBc (Core) protein, the unique capsid component. Once in the nucleus, the rcDNA is repaired by cellular factors, chromatinized, and loaded with cellular and viral proteins to form cccDNA, the viral double-stranded DNA episome responsible for the production of viral RNAs [7]. This 3.2 Kb viral genome contains 2 enhancers (enhancer 1 and 2) and 4 promoters (preCore/Core, preS1, preS2, and X) that harbor binding sites for several ubiquitous and hepatocyte-specific transcription factors, and drive the transcription of 5 unspliced and several spliced RNAs [8–10]. Seven viral proteins are translated from unspliced transcripts and are sufficient to initiate productive replication. Among them, HBx is a 154 amino-acid protein translated from mRNAs produced from enhancer 1/X promoter with diverse transcription initiation sites [11]. This viral protein is absolutely required to initiate and maintain cccDNA in a transcriptionally active state [12,13]. The reason for this requirement remained obscure for a long time until the discovery that, in the absence of HBx, cccDNA is bound and silenced by the Smc5/6 cohesin complex. The transcriptional repression of cccDNA is relieved by HBx by hijacking the CUL4A E3 ubiquin ligase to induce the degradation of the Smc5/6 complex [14–16].

In addition to HBx, the viral HBc protein, translated from the pregenomic RNA (pgRNA) transcribed from the enhancer 2/preCore/Core promoter, is also suspected of playing important regulatory roles in the production of viral RNAs [17]. Indeed, besides its essential role to form capsids in the cytoplasm, HBc is also present in the nucleus where, as HBx, it associates with cccDNA [18,19]. The consequences of this interaction on viral transcription are, however, still a matter of debate [20–23]. The recent observations that capsid-derived HBc stably associates with the viral genome once released into the nucleus [24], and that it interacts with Hira, a histone variant deposited on cccDNA during the chromatinization process [25], additionally suggests that it may be involved during cccDNA repair and/or chromatinization.

In a previous study, we found that HBc interacts with several cellular RBPs within the hepatocyte’s nucleus raising the hypothesis that it may hijack their activities to control viral replication [26]. In the same study, we discovered that a drug named 1C8, capable of inhibiting the phosphorylation of some RBPs associated with HBc, could reduce the level of nascent HBV RNAs [26]. These results suggested that one or more protein kinases targeted by this 1C8 might control the synthesis of viral RNAs by phosphorylating cellular and/or viral proteins.

The objective of the current study was to identify the kinase(s) targeted by 1C8 responsible for its (their) effect on cccDNA transcriptional activity. Our results pointed to DYRK1A as one of these kinases. DYRK1A is a pleiotropic serine/threonine kinase that, among other activities, regulates the transcription of cellular genes by associating with chromatin and phosphorylating histones, chromatin regulators as well as the carboxy-terminal domain of the RNA polymerase II (RNA Pol II) [27–29]. DYRK1A also participates in nuclear condensates associated to transcription elongation [30]. We show that DYRK1A interacts with cccDNA and up-regulates the transcription of viral RNAs. In addition, our data strongly suggest that DYRK1A transcriptional activity is mediated by the recognition of a consensus motif within enhancer I/X promoter region that drives HBx production. Therefore, we propose that DYRK1A is a novel cellular proviral factor that may participate HBV life cycle by stimulating the transcription of HBx, a viral factor required to unleash the complete HBV transcriptional program leading to productive replication.

## MATERIALS AND METHODS

### Cell culture and infection

HepaRG cells were cultured, differentiated, and infected by HBV as previously described [31]. Primary human hepatocytes (PHHs) were freshly prepared from human liver resection obtained from the Centre Léon Bérard and Hôpital Lyon Sud (Lyon) with French ministerial authorizations (AC 2013-1871, DC 2013 – 1870, AFNOR NF 96 900 sept 2011) as previously described [32]. To generate HepaRG cells lines over-expressing HA-tagged human DYRK1A and DYRK1B (wild-type and kinase-dead versions (KR), in which the ATP binding Lys179 was replaced by Arg),) and FLAG-tagged SRPK1 [33], the open reading frame of each kinase was inserted into the pLenti4/TO lentiviral vector plasmid, under the control of the minimal CMV/TetOn promoter. Lentiviral particles generated from these plasmids were used to transduce HepaRG-TR cells [13]. Transduced cells were selected using blasticidin (10 μg/mL) and zeocyn (100 μg/mL), amplified and frozen as polyclonal lines. HuH7-NTCP cells were cultivated in 10% fetal calf serum supplemented DMEM (4.5 g/L glucose) [34]. HBV genotype D inoculum (ayw subtype) was prepared from HepAD38 [35] cell supernatant by polyethylene-glycol-MW-8000 (PEG8000, Sigma) precipitation (8% final) as previously described [36]. The titer of endotoxin free viral stocks was determined by qPCR. Cells were infected overnight in a media supplemented with 4% final of PEG, at indicated multiplicity of infections (MOIs), as previously described [31]. Measure of secreted HBs and HBe antigens was performed by chemo-luminescent immune assay (AutoBio, China), following manufacturer’s instructions and expressed as international units/mL (IU/mL) and Paul Erlich Internation units/mL (PEIU/mL), respectively.

### Chemical compounds

Unless otherwise specified, chemical reagents, drugs, antibiotics were purchased from Sigma Aldrich. Tenofovir (TFV) was a kind gift of Gilead Sciences (Foster city, USA). The core assembly modulator (CAM) used in the experiments was previously described [37], and resynthesized by AI-Biopharma (Montpellier, France). 1C8, *i.e.* 1-Methyl-N-(5-nitrobenzo[d]isothiazol-3-yl)-4-oxo-1,4-dihydropyridine-3-carboxamide, was synthesized and purified at 99% by AGV (Montpellier, France). RG7834 [38], a molecule that destabilizes HBV RNAs, was synthetized by AI-Biopharma (Montpellier, France). AZ191 was purchased from SelleckChem, IFN-α (Roferon) from Roche, SRPIN340 and TG003 from MedChem Express. DB18 was a kind gift from René Grée (University of Rennes, France) ([39].

### siRNA Transfection

Differentiated HepaRG (dHepaRG) cells or PHHs seeded into a 24-well plate were transfected with 25 nM of siRNA using Lipofectamine RNAiMax (Life Technologies), following manufacturer’s instructions: siDYRK1A (Dharmacon SmartPool L-004805-00), siDYRK1B (Dharmacon SmartPool L-004806-00), siCLK2 (Dharmacon J-004801-10), siSRPK1 (Dharmacon SmartPool: L-003982-00) and siControl (siCTL) (Dharmacon D-001810).

### Viability/cytoxicity assays

Neutral red uptake and sulforhodamine B assays were performed to estimate cell viability/cytotoxicity as previously described [40]. Puromycin (Puro), was used as a positive control at a concentration of 10 µM.

### Nucleic acid extraction and analysis

HBV RNAs and DNA were extracted from cells with the Nucleospin RNA (Macherey-Nagel) and MasterPureTM Complexe Purification Kit (BioSearch Technologies) kit without digestion with proteinase K [41], respectively, according to the manufacturer’s instruction. RNA reverse transcription was performed using Maxima First Strand cDNA Synthesis kit (Thermofisher). Quantitative PCR for HBV were performed using HBV specific primers and normalized to PRNP or RPLP0 housekeeping genes as previously described [42]. Pregenomic RNA (pgRNA) was quantified using the TaqMan Fast Advanced Master Mix (Life Technologies) and normalized to GUS B cDNA levels. HBV cccDNA was quantified from total DNA by TaqMan qPCR analyses and normalized to β-globin DNA level, as previously described [43]. Unless otherwise stated, results are expressed as the mean normalized ratio +/- SD, of 3 independent experiments, each performed in triplicate.

### Western blot analyses

Proteins were resolved by SDS-PAGE and then transferred onto a nitrocellulose membrane. Membranes were incubated with the primary antibodies corresponding to the indicated proteins. Proteins were revealed by chemiluminescence (Super Signal West Dura Substrate, Pierce) using a secondary peroxydase-conjugated antibody (Dako) at a dilution of 1:10000. Primary antibodies used were CLK1 (Clinisciences, ARP52021, 1:1000), CLK2 (Abcam, Ab65082, 1:1000), SRPK1 (Abcam, Ab131160, 1:1000), DYRK1A (Abcam Ab69811, 1:1000 and SantaCruz sc-100376, 1:1000), DYRK1B (Cell Signaling 5672, 1:1000), HBc (provided by A. Zlotnick (Indiana University, USA), 1:40000), β-tubulin (Abcam 6046,1:10000), β-actin (Sigma A2228, 1:2000), cyclin D1 (Cell Signaling 55506, 1:2000), HA (Abcam 130275; 1:1000), and FLAG (Sigma F1804; 1:1000).

### Native cccDNA-immunoprecipitation (cccDNA-IP) and ChIP

cccDNA was immunoprecipitated from frozen cell pellets following a modified native chromatin-immuno-precipitation (ChIP) protocol as previously described [24]. Antibodies (2 µg) used were: DYRK1A (Abcam 69811 and Sigma D1694), HBc (Life Invitrogen MA1-7609), total RNA Pol II (Diagenode C15200004), RNA Pol II phosphor-Ser5 (Abcam Ab5131), RNA Pol II phosphor-Ser2 (Abcam Ab5095) and control IgG (Cell Signaling 2729).

### Luciferase assays

Huh7.NTCP cells seeded in a 48-well plates were transfected using TransIT-LT1 transfection reagent (Mirus) and lysed 72 hours later using the Bright-Glo Luciferase reagent (Promega), according to the manufacturer’s instructions. Luciferase activity was quantified on a Luminoskan (Life Technologies). Plasmids expressing luciferase under the control of HBV promoters were purchased from Addgene: preC/C_Luc (pHBV_Luc; ref:71414); pX_Luc (pHBV-X/EnhI-Luc; ref:71418). Mutations M1 (<u>CAGGTG</u>cAac) and M2 (<u>CAGGTGTACA)</u> in pX_Luc were introduced using Q5 site-directed mutagenesis kit (New England Biolabs). Plasmids expressing DYRK1A or DYRK1A_KR under the control of the CMV promoter were previously described [29].

### Run-ON

HBV nascent RNAs were quantified as previously described [44,45]. Briefly, cells were incubated for 2h at 37°C with 5-bromouridine (BrU, Sigma), before RNA extraction and immunoprecipitation using an anti-BrdU antibody (BD Pharmingen) and Dynabeads coated with anti-mouse IgG (Life Technologies), overnight at 4°C. Captured RNAs were then eluted, concentrated and quantified by RT-qPCR. Total RNAs, extracted before IP were similarly quantified by RT-qPCR. Controls were provided by cells treated with actinomycin D (Invitrogen, 1 µM; 1 h before labeling), RG7834 (0.05 µM, added twice 5 and 3 days before labeling) or cells incubated in the absence of BrU.

### Statistical analysis

Statistical analyses were performed using the GraphPad Prism 9 software and a two-tailed Mann-Whitney non-parametric tests. In graph bars, error bars represent standard deviation (SD) unless otherwise indicated. A p value ≤ 0.05 was considered as significant: *, p value ≤ 0.05; **, p value ≤ 0.01; ***, p value ≤ 0.001.

## RESULTS

### 1C8 targets five CMGC kinases expressed in human hepatocytes

Our previous results indicated that the compound 1C8, an inhibitor of the phosphorylation of some cellular RBPs [46,47], downregulated the synthesis of HBV RNAs without affecting cccDNA levels [26]. This observation suggested that one or more kinases involved in RBPs phosphorylation had a proviral function on HBV transcription. A kinase profiler assay conducted using 1C8 (Eurofins Pharma Discovery) identified 18 different kinases highly inhibited by the compound (Figure S1) [48]. Among them, the members of the Cdc2-like kinases (CLK) family CLK1, CLK2 and CLK4, and those of the dual-specificity tyrosine phosphorylation-regulated kinases (DYRKs) family DYRK1A and DYRK1B, were found. All these kinases that belong to the CMGC group of the eukaryotic kinome [49,50], phosphorylate RBPs, in particular SR proteins, and their inhibition has been proposed to have therapeutic potential [51,52]. Interestingly, the serine-arginine protein kinases (SPRKs) family that also belongs to the CMGC group and are well-known regulators of SR proteins, were not found as 1C8 targets.

One first requirement for the involvement of these kinases in HBV life cycle is their expression in hepatocytes. Thus, we analyzed the level of expression of these kinases in the human hepatocyte cell line HepaRG, using either proliferating or differentiated (dHepaRG) cells, and in primary human hepatocytes (PHHs), whether infected or not with HBV. We detected the expression of CLK1 and CLK2 mRNAs (CLK4 was not measured as it is highly similar to CLK1) in HepaRG cells, irrespectively of their differentiation or infectious status. In comparison, the mRNA levels of DYRK1A and DYRK1B were much lower (Figure S2A). Similar results were observed in PHHs and in human liver biopsies (Figure S2B and C). Moreover, the four kinases were detected at the protein level in both cell types (Figure 1). We conclude that the four kinases targeted by 1C8 are expressed in human hepatocytes.

**Figure 1.**
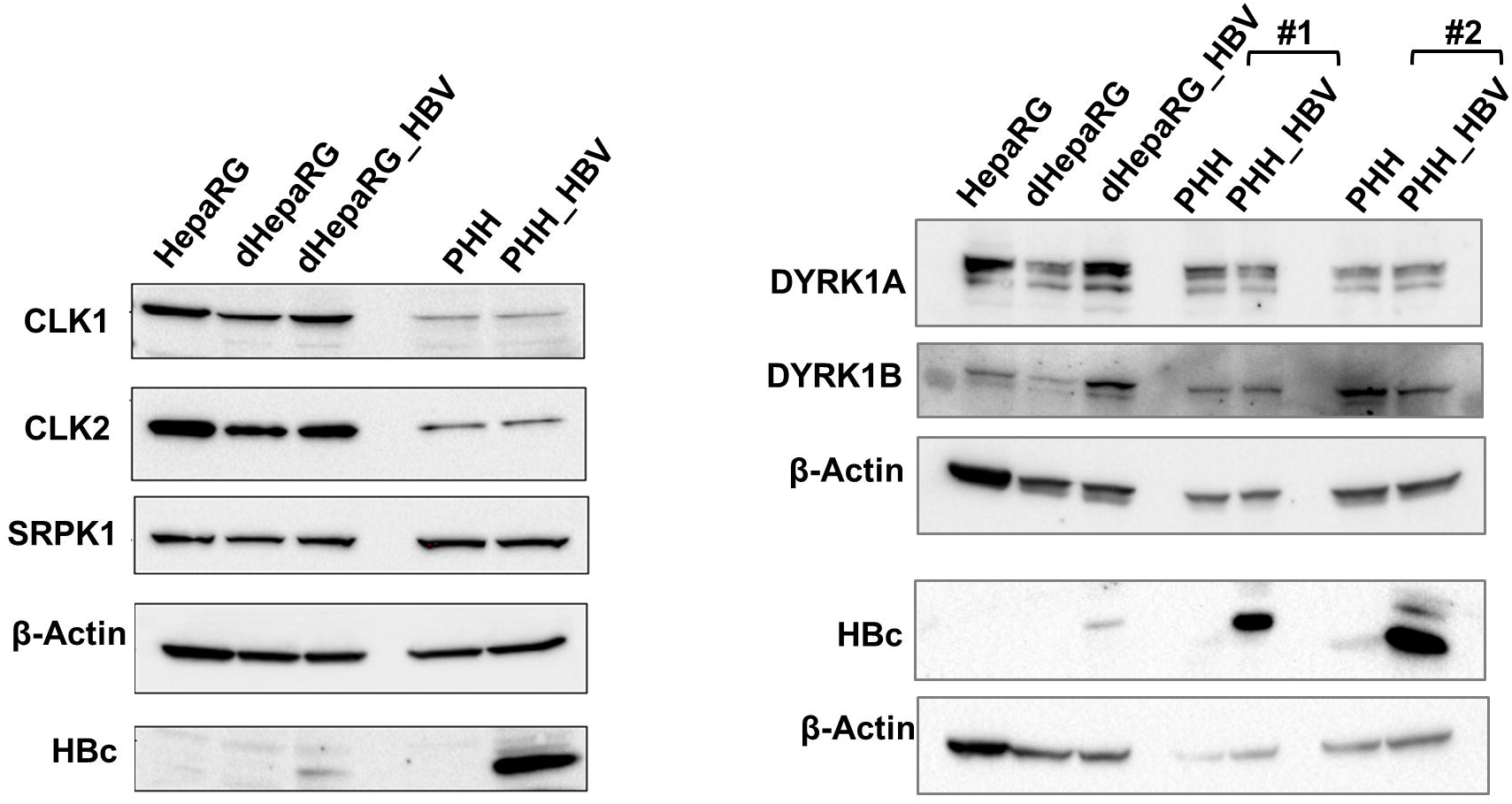
Expression levels of selected CMGC kinases in human hepatocytes. Western blot analysis of CLK1, CLK2, SRPK1, DYRK1A, DYRK1B expression in human hepatocytes (HepaRG cells or PHHs), mock- or HBV-infected. HepaRG: non-differentiated cells; dHepaRG: differentiated cells; dHepaRG_HBV: HBV-infected dHepaRG; PHH: mock-infected cells; PHH-HBV: HBV-infected cells. #1 and #2 refers to PHH purified from two different donors. The detection of HBc is used as a control for HBV infection. β-Actin is used as loading control.

### Inhibitors targeting DYRK1A/B, but not CLK kinases, reduce HBV replication

In order to identify which CLK/DYRK kinase(s) targeted by 1C8 potentially regulates HBV replication, the effect of inhibitors of these kinases was analyzed on the production of HBV RNAs and the secretion of HBs and HBe antigens in HBV-infected PHHs, two traits commonly used to measure HBV productive infection (Figure 2). As positive controls, we used IFN-α, an inhibitor of HBV transcription [53], which reduces HBV RNAs, and a class I CAM, which prevents capsid assembly [54], and also reduces HBe secretion (Figure 2B) [37]. Treatment of cells with either SRPIN340 [55], an inhibitor of the kinase SRPK1, not targeted by 1C8, or with TG003, a well-known CLK inhibitor [56], had no effect on HBV RNA production or HBs and HBe secretion (Figure 2B).[55] The absence of effect of inhibitors targeting CLKs was further confirmed using DB18 (Figure S3), a compound that selectively targets CLK1/4 and CLK2 but not DYRK1A [39]. In contrast, both 1C8 and AZ191, a DYRK1 kinases inhibitor but with no effect on CLKs [57], induced a decrease in HBV RNA levels (Figure 2B and S3). The effect of AZ191 was stronger than that of 1C8 (Figure 2B), with no evidence of cell toxicity at the dose of 10-20 µM used (Figure S4). The lower effect of 1C8 could be due to the poor solubility of 1C8 and/or to a stronger susceptibility to the hepatocyte detoxifying activity. Altogether, these results pointed towards DYRK1A/1B as potential kinases able to directly or indirectly regulate HBV replication.

**Figure 2.**
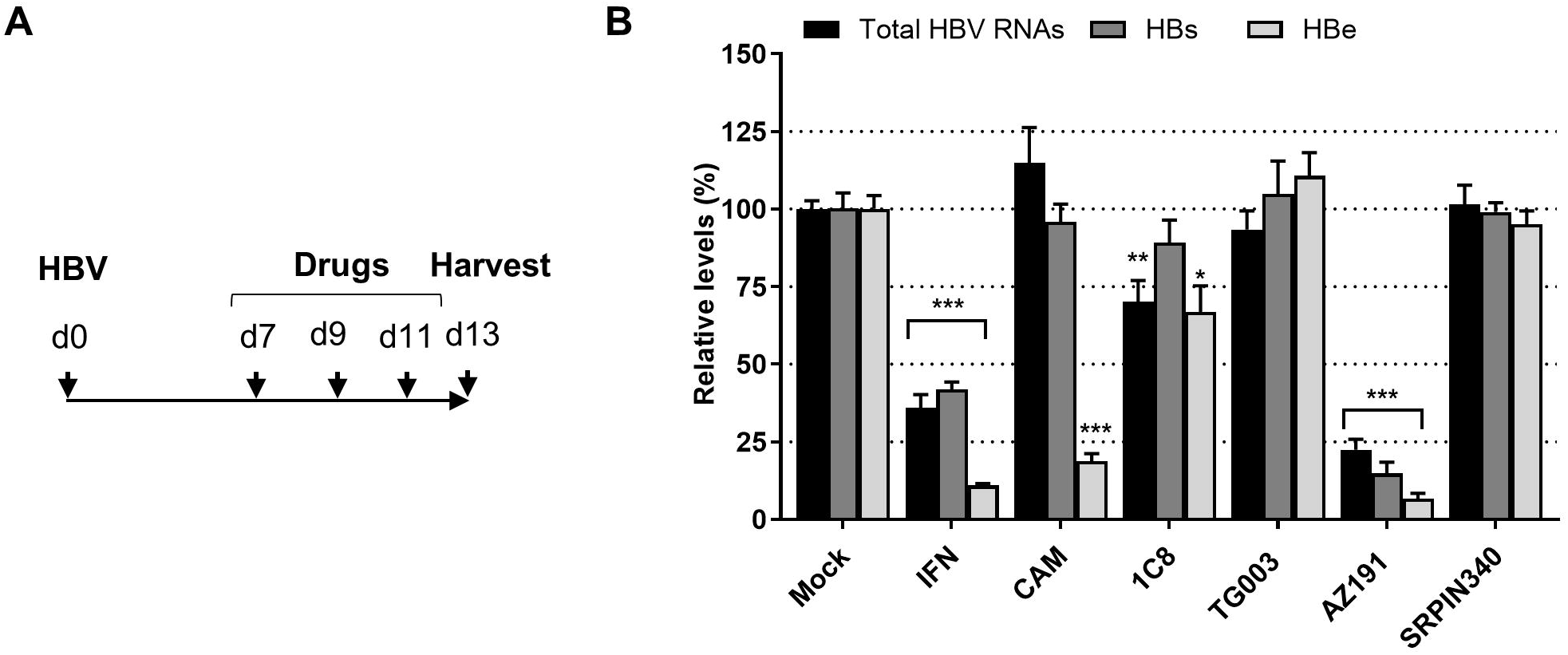
Chemical inhibitors targeting DYRK1A and DYRK1B reduce HBV RNAs production. **A.** Experimental outline. PHHs derived from different donors were infected with HBV (MOI: 500 vge/cell) and seven days later, treated three times with the indicated compounds. Viral parameters were measured 2 days after the last treatment. IFN: Interferon-α (500 IU/mL), CAM: capsid assembly modulator (class I, 10 µM), 1C8 TG003, AZ191, and SRPIN340 were used at 20 µM. **B.** Relative quantification of total intracellular viral RNAs, and of secreted HBs and HBe antigen levels, as indicated. Statistical values are from comparisons between the treated and mock-treated cells for each of the measurements and expressed as the mean +/- SD of 3 independent experiments, each performed in triplicate.

### Production of HBV RNAs responds to DYRK1A/DYRK1B levels

To provide further support to the results with the kinase inhibitors, we set up knock-down (KD) experiments in HBV-infected dHepaRG cells, using a well-established protocol in our lab, consisting in two successive siRNAs transfections followed by downstream analyses 10 days post-transfection (dpt) (Figure S5A). Unfortunately, despite several attempts using different sources of siRNA, efficient KD of CLK1 mRNA was never achieved in hepatocytes, precluding further analysis with this kinase. It is possible that a previously described stress-responsive mechanism, able to rapidly compensate for the loss of CLK1 [58], may be the cause of this failure [58]. In contrast, CLK2 and SRPK1 expression was reduced in HBV-infected dHepaRG cells upon siRNA treatment, but no effect on HBV RNAs was observed (Figure S5B and C). Transfection of siRNAs targeting either DYRK1A or DYRK1B resulted in the visible death of most cells between 7- and 10-dpt (Figure S6). This cytopathic effect led us to modify the depletion protocol to investigate effects at earlier time-points (Figure S7A), during which no toxic effects were observed (Figure S7B and C), but with KD efficiency gradually increasing (Figure S7D). In these conditions, total viral RNA levels remained unaffected when targeting DYRK1B, but they were clearly reduced when targeting DYRK1A (Figure S7E). Importantly, DYRK1A KD induced not only a significant decrease in total HBV RNAs, but also in pgRNA and downstream parameters (HBe and HBs secretion), without affecting cccDNA levels (Figure 3).

**Figure 3.**
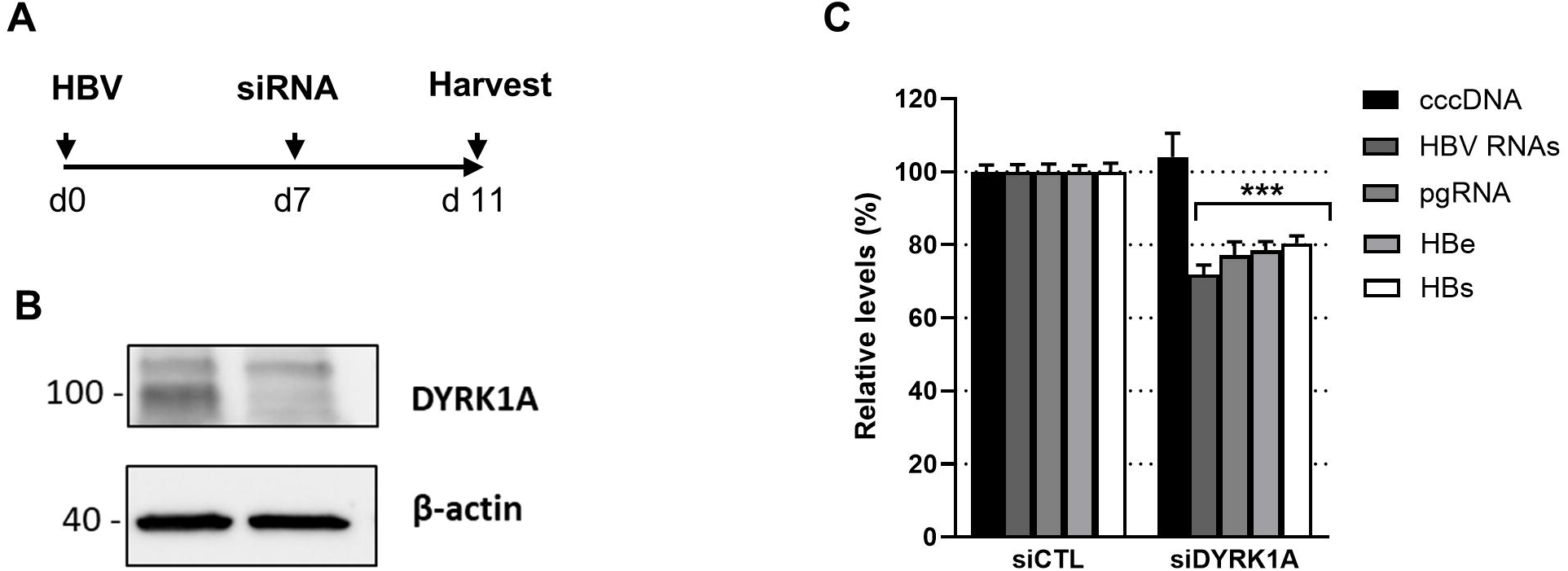
Knock-down of DYRK1A decreases HBV RNAs levels in HBV-infected dHepaRG cells. **A.** Experimental outline. Cells were infected and 7 days later transfected with siRNAs targeting DYRK1A. DYRK1A protein levels (**B**) and the viral parameters indicated (**C**) were quantified 4-dpt, a time point at which cell toxicity was not observed (Figure S6). Statistical values are from comparisons between siDYRK1A and siCTL for each of the measurements and expressed as the mean +/- SD of 3 independent experiments, each performed in triplicate.

To confirm this result, the effect of DYRK1A KD was also assessed in HBV-infected PHHs, in which the effect of DYRK1A KD could be evaluated at longer times after transfection due to the lack of toxicity of the siRNA treatment with good depletion efficiency (Figure S8A and B). Surprisingly, discordant results were obtained using this cell type. For 3 out of 7 total experiments, each performed with PHHs from independent donors, KD of DYRK1A resulted in a decrease of viral RNAs without affecting cccDNA levels (Figure S8C and D), in agreement with the results in dHepaRG cells. However, in four of these experiments, the opposite result was observed despite similar levels of DYRK1A KD (Figure S8E and F). One possibility for these discordant results could be related to compensatory effects between DYRK1A and its close paralog DYRK1B [49,50], effects that might have variable penetrance given that PHHs are from different donors. In fact, an increase in DYRK1B levels in response to silencing DYRK1A has been shown in a different physiological context [59]. To test this possibility, we quantified the levels of DYRK1A and DYRK1B mRNAs in control and KD cells by RT-qPCR. The analyses revealed that DYRK1B mRNA was up-regulated in PHHs transfected with siRNA against DYRK1A. In addition, the up-regulation of DYRK1B mRNA was significantly higher in PHHs where HBV RNAs increased following DYRK1A KD, compared to PHHs where the opposite effect was observed (Figure S9A), suggesting a possible positive correlation between the level of DYRK1B compensation and the effect of DYRK1A KD on HBV RNAs. The increase in DYRK1B upon DYRK1A KD was also observed in dHepaRG cells but at a relatively lower level compared to PHHs, despite a similar efficiency in DYRK1A KD (Figure S9B), highlighting an additional layer of complexity linked to the cell line used. In contrast, DYRK1A mRNA levels were not upregulated upon DYRK1B KD in either dHepaRG cells or PHH (Figure S9B and S10B). Importantly, the variations in DYRK1B mRNA were confirmed at the protein level (Figure S10A). Interestingly, in those PHHs where DYRK1B compensation upon DYRK1A KD was higher, DYRK1B KD, either alone or in combination with DYRK1A, induced a decrease in HBV RNA (Figure S10C). Altogether, these results indicated a regulatory cross-talk between DYRK1B and DYRK1A whereby DYRK1A depletion is compensated by an increase in DYRK1B mRNA, either due to the existence of DYRK1A-responsive elements within the DYRK1B promoter region or to DYRK1A-dependent cellular events that impact on DYRK1B transcription and/or mRNA stability. More importantly, the level of DYRK1B compensation provided a rational explanation for the divergent results observed upon DYRK1A KD in PHHs.

Next, we wondered whether DYRK1A could stimulate HBV RNAs when overexpressed. To this end, HepaRG cells over-expressing either a wild-type (wt) DYRK1A or a kinase-dead version (DYRK1A_KR), under the control of a tetracyclin (Tet)-inducible promoter were developed (Figure 4A and B). Notably, over-expression of DYRK1A_WT, but not of SRPK1, led to an increase in HBV parameters, either intracellular viral RNAs or extracellular HBs and HBe antigens (Figure 4C-F). This effect was partially dependent on DYRK1A catalytic activity, as overexpression of the kinase-dead version induced a much lower increase (Figure 4C-F), further supporting the kinase activity dependence showed by the experiments with the inhibitors. Interestingly, overexpression of DYRK1B induced only a modest increase in HBV pgRNA but not in total RNAs, indicating a lower capacity to modulate viral RNAs compared to DYRK1A (Figure S11).

**Figure 4.**
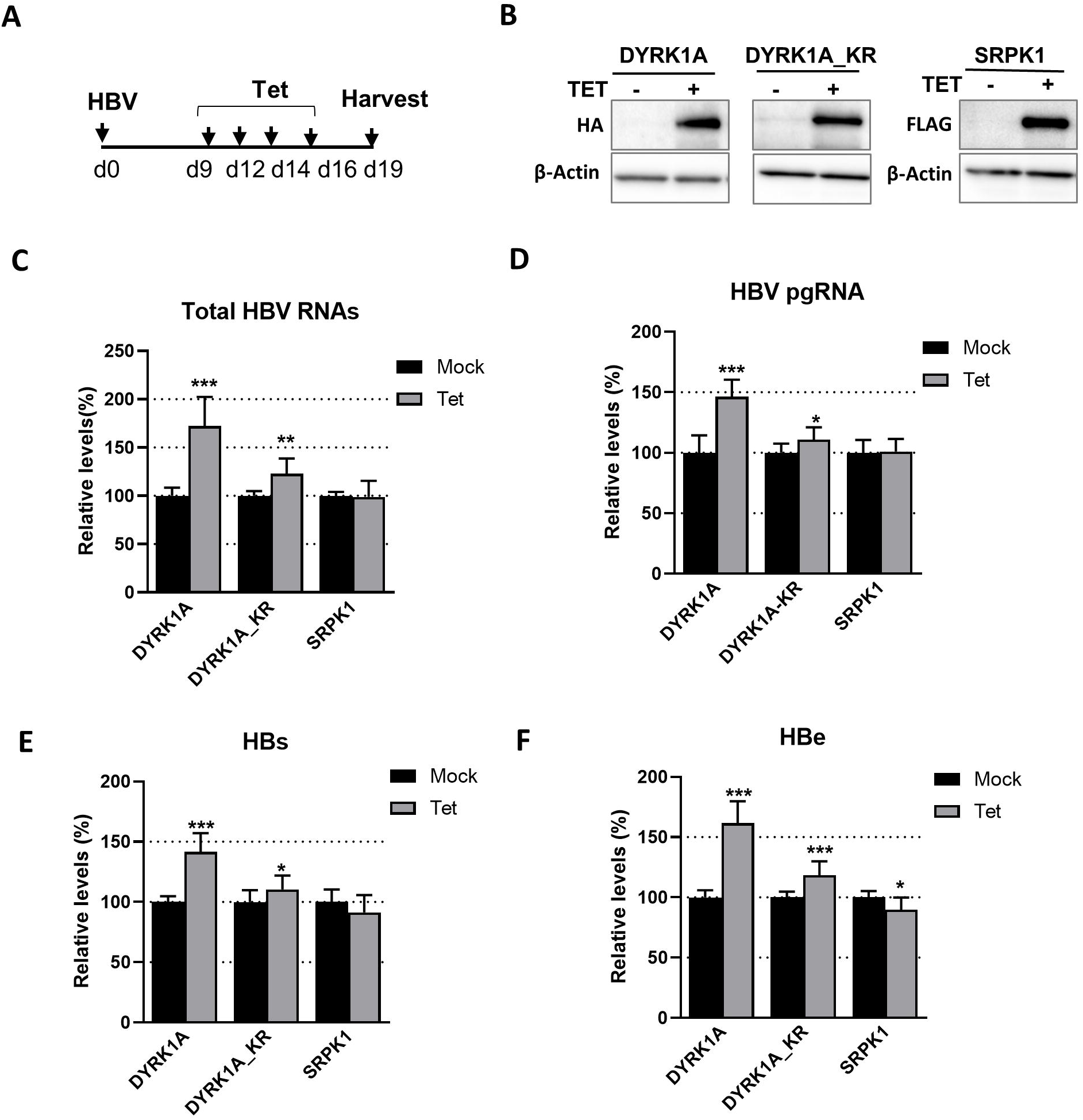
Over-expression of DYRK1A increases HBV RNA levels in HBV-infected dHepaRG cells. **A**. Experimental outline. dHepaRG cells engineered to express either HA- DYRK1A, HA-DYRK1A_KR or FLAG-SRPK1, under the control of a tetracyclin (Tet)-inducible promoter, were infected with HBV (500 vge/cell) with additions of Tet (5 µg/mL) or mock-treated every 2-3 days for ten days. Western blot analyses were performed to detect the over-expressed kinases with anti-tag antibodies (**B**) and viral parameters (**C** to **F**). Results are normalized to the mock situation and expressed as the mean +/- SD of 4 independent experiments, each performed in triplicate.

Altogether, the loss-of-function and gain-of-function studies indicate that DYRK1A could exert a proviral function by increasing the levels of HBV RNAs, and that its catalytic activity is involved in this function. They also highlight a complex cell-specific compensatory mechanism between DYRK1A and DYRK1B expression, and, at least partially, redundant functions on HBV replication.

### DYRK1A associates to HBV cccDNA and up-regulates transcriptional activity from enhancer 1/HBx promoter region

Previous studies indicated that DYRK1A promotes the transcription of a set of cellular genes by associating to chromatin at RNA Pol II promoter regions and close to transcriptional start sites (TSS) [29]. We therefore investigated whether DYRK1A could associate to HBV cccDNA using our recently optimized native ChIP protocol [24]. This protocol is efficient in detecting the presence of HBc bound to cccDNA (Figure 5A), and showed that DYRK1A was associated to cccDNA in HBV-infected dHepaRG cells (Figure 5A). The same result was obtained in HBV-infected PHHs with two different antibodies to DYRK1A (Figure 5B and C). These results indicated that DYRK1A interacts with cccDNA, either directly or indirectly via other cellular and/or viral factors.

**Figure 5.**
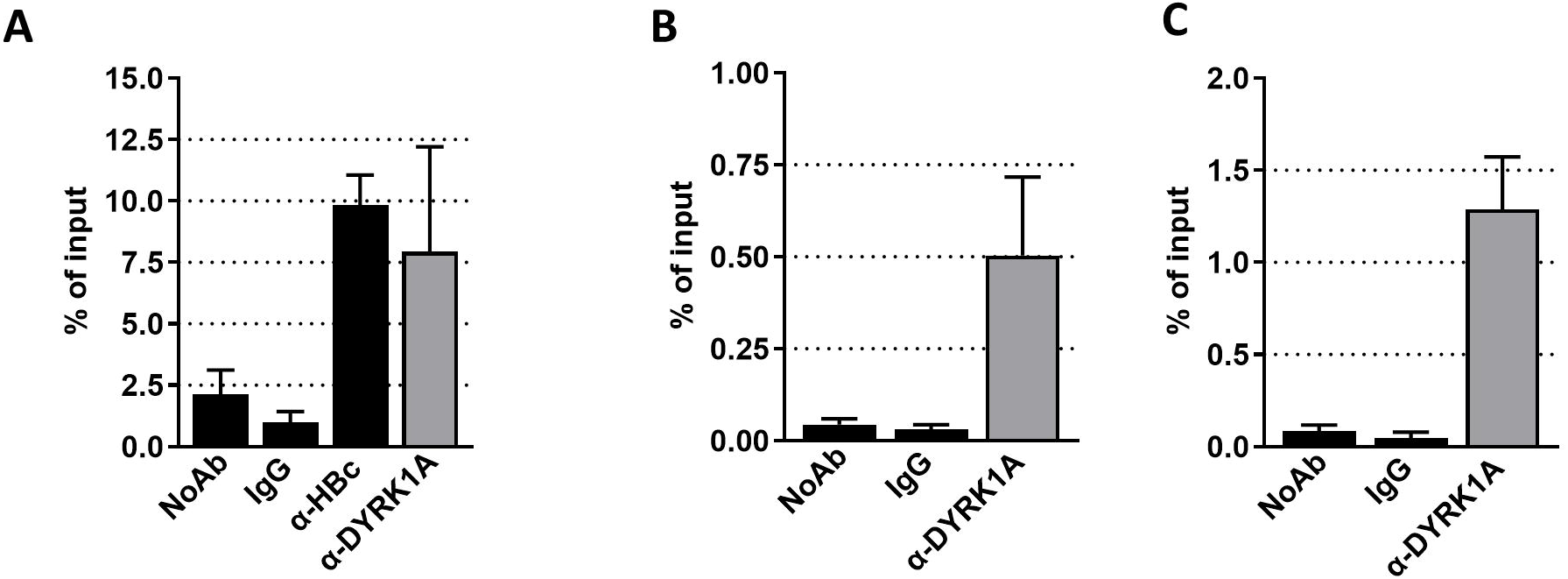
DYRK1A interacts with HBV cccDNA. Native ChIP analyses were performed on HBV-infected dHepaRG (**A**) or PHH (**B** and **C**) using antibodies against HBc or DYRK1A. The results obtained with two different antibodies against DYRK1A are shown in panels **B** and **C**. cccDNA was quantified from immuno-precipitated material by TaqMan qPCR. Results represent the mean +/- the standard error of the mean of 3 (dHepaRG) or 5 (PHHs) independent experiments.

DYRK1A recruitment to chromatin is mediated by a 10 nucleotides sequence TCTCGCG(A/G)(G/T)(A/G) located near TSSs [29,60] (Figure 6A). Association of DYRK1A to such sites regulates the transcription of downstream genes by phosphorylating the C-terminal domain (CTD) of RNA Pol II [29]. Interestingly, a sequence partially matching the DYRK1A motif is present in the enhancer 1/X promoter region of the HBV genome, upstream of the HBx TSSs (Figure 6A). To investigate whether the enhancer 1/X promoter could be a target of DYRK1A, reporter assays were conducted with constructs containing the luciferase cDNA under the control of either the enhancer 1/X promoter (pX_Luc) regions or enhancer 2/preCore/Core (preC/C_Luc) as a negative control, co-transfected together with constructs expressing either DYRK1A wt or DYRK1A-KR. As shown in Figure 6B, DYRK1A expression significantly increased the activity of the enhancer 1/X promoter reporter, but not that of the enhancer 2/preCore/Core reporter. Interestingly, a lower but still significant enhancement was observed following transection of the plasmid expressing the DYRK1A kinase-dead version. In this regard, it is worth noting that a transcriptional effect independent of the DYRK1A kinase activity was observed in one-hybrid assays [29], suggesting a possible scaffolding activity. To further determine if the DYRK1A transactivating activity was mediated by the putative DYRK1A motif within enhancer 1/X promoter, two different mutant motifs were generated (Figure 6A). Both pX_M1_Luc and pX_M2_Luc constructs no longer responded to DYRK1A (Figure 6C), indicating that the enhancing activity of DYRK1A did depend on the presence of the motif characteristic of its recruitment to cellular chromatin. Altogether, these results indicated that DYRK1A associates to cccDNA and strongly suggested that its effect on HBV RNAs was, at least in part, linked to its capacity to recognize a sequence matching the DYRK1A motif present on the enhancer 1/HBx promoter region.

**Figure 6.**
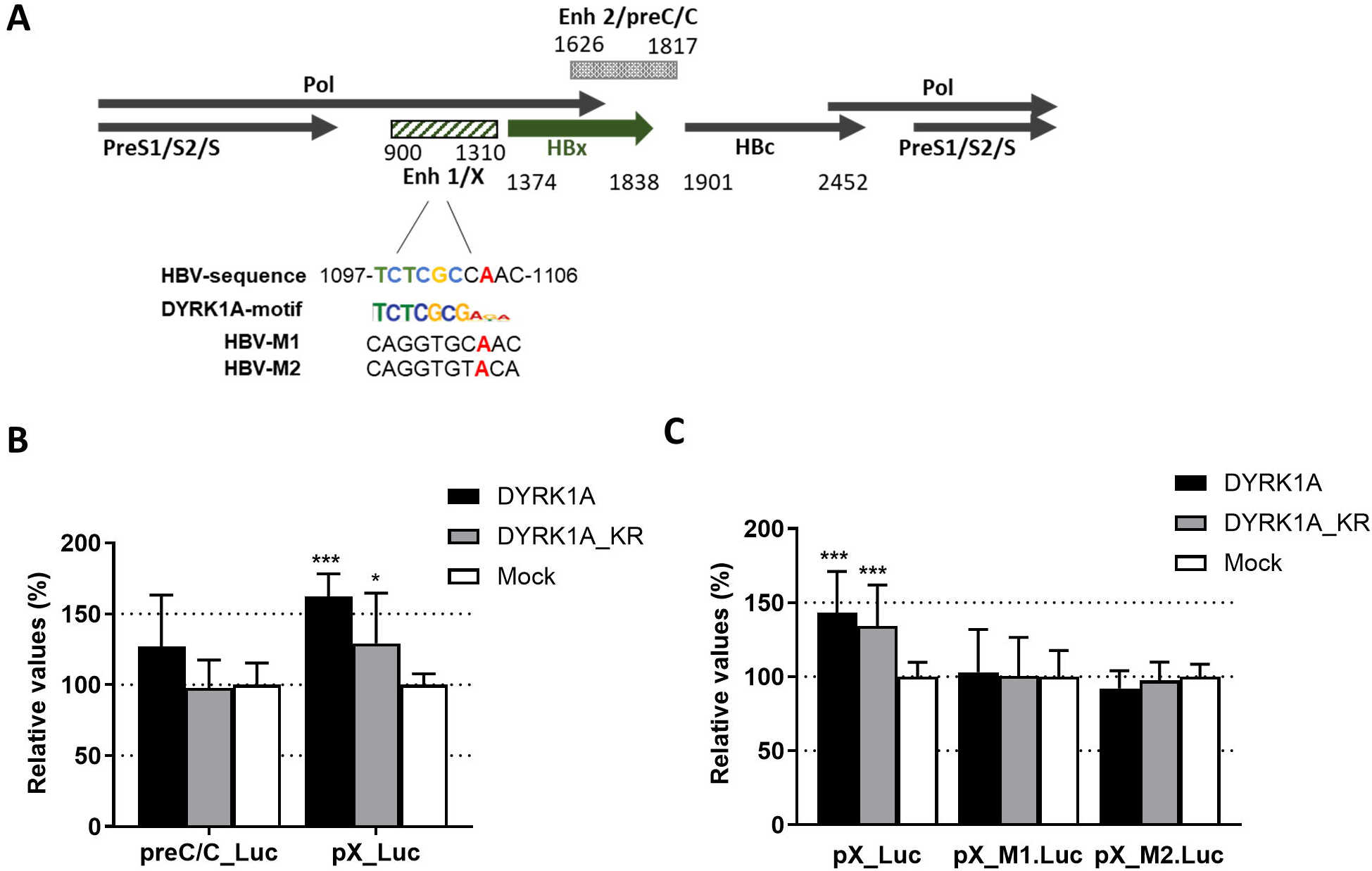
DYRK1A activity on HBV transcription regulatory regions. **A.** Schematic view of a linearized HBV genome (genotype D, ayw subtype), showing the position of the four main ORFs (the HBx ORF is indicated by a large green arrow) and of the two enhancer/promoter regions. The enhancer 1/X promoter region contains a putative DYRK1A recognition site that partially matches the consensus motif found in DYRK1A responsive cellular promoters. The two mutation M1 and M2 introduced within enhancer 1/X promoter are shown below. Nucleotide positions are numbered from the unique EcoRI site. **B.** Plasmid containing the luciferase cDNA under the control of the enhancer 1/HBx (pX_Luc) promoter or the enhancer 2/preCore/Core (preC/C-Luc) regions were cotransfected together with a plasmid coding for DYRK1A, DYRK1A_KR, or a control plasmid into Huh7-NTCP cells. Luciferase assays were performed three days later. **C.** Luciferase assays were performed by transfecting plasmids containing either a wt or two different mutated putative DYRK1A motifs. Results are normalized to the cells co-transfected with a control plasmid (Mock) and expressed as the mean +/- SD of 4 independent experiments, each performed in triplicate.

### Over-expression of DYRK1A up-regulates the transcription of HBV RNAs

To investigate whether DYRK1A acts at the transcriptional level, RUN-On assays were performed. For this, BrU-labeled RNAs were extracted from HBV-infected dHepaRG cells, with DYRK1A (WT or KR versions) expression induced by Tet treatment. To proof that the experimental set up does detect nascent RNAs, we show that treatment of cells with RG7834, a compound that degrades HBV RNAs [38,44], had no effect on nascent RNAs, whereas actinomycin D, an inhibitor of RNA Pol II, blocked their synthesis (Figure 7). As shown above, over-expression of DYRK1A wt increased total HBV RNAs, whereas its kinase inactive version had no effect (Figure 7, left panel). Importantly, up-regulation of nascent HBV RNAs was observed only when the expression of active DYRK1A was induced (Figure 7, right panel), strongly suggesting that the DYRK1A-dependent effect takes place at the transcriptional level.

**Figure 7.**
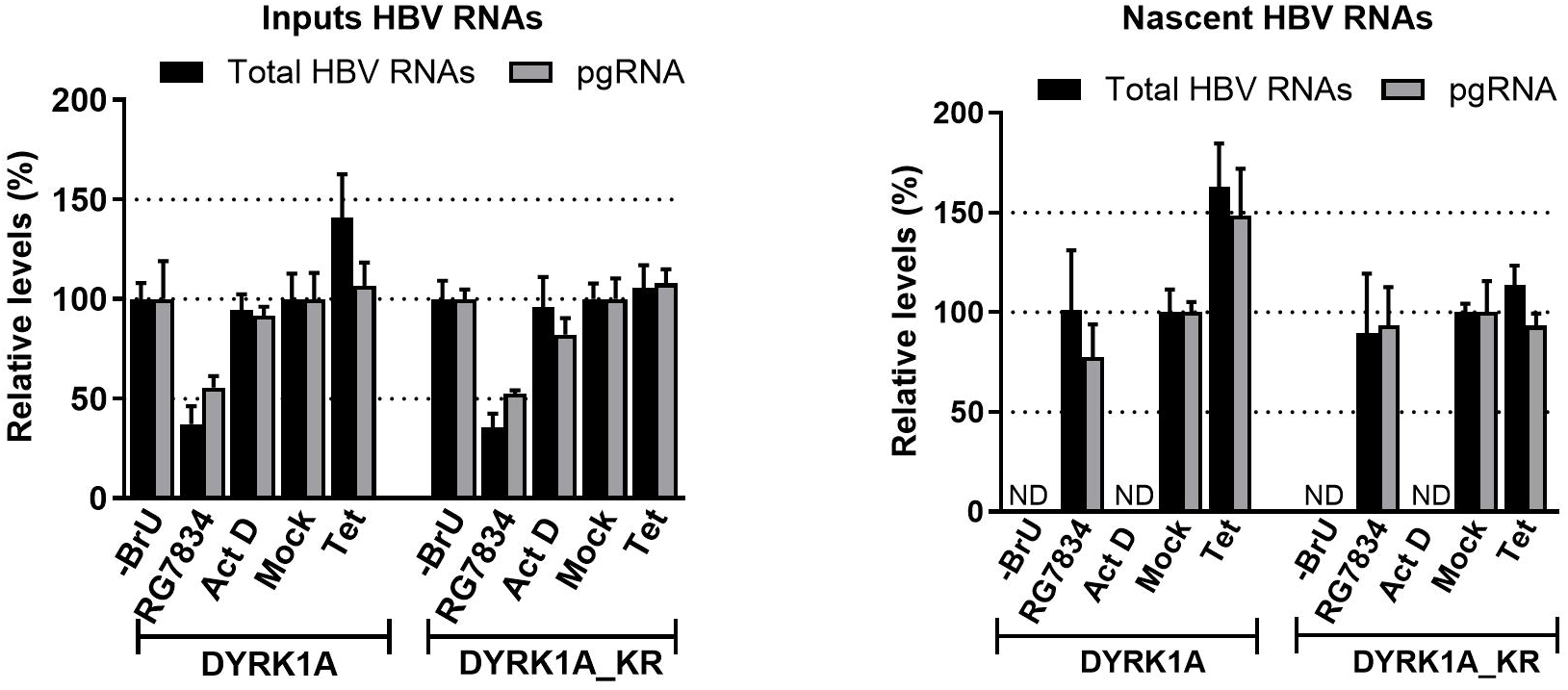
DYRK1A over-expression increases the level of HBV nascent RNAs. Run-On analysis. dHepaRG-TR-DYRK1A or -DYRK1A_KR cells were infected with HBV and treated with Tet or mock-treated as indicated in the legend of Figure 6A. Controls were provided by cells incubated in the absence of BrU (-BrU) or in the presence of actinomycin D (Act D) or RG7834. Labeled RNAs were immunoprecipitated using an anti-BrdU antibody. Input HBV RNAs (left panel) and immunoprecipitated RNAs representing nascent RNAs (right panel) were quantified by RT-qPCR. Results are expressed as the mean+/- SD of 2 independent experiments, each performed in triplicate. ND: value under the detection limit.

## DISCUSSION

So far, only few cellular kinases were reported to modulate the HBV life cycle. In addition, most if not all of them, exert their effects at late steps of the infectious cycle, during capsid assembly and pgRNA packaging [17]. This is the case of CDK2, PKC and PLK1, which phosphorylate the HBc protein during capsid assembly in the cytoplasm [61–64]. In this study, we asked whether cellular kinases could also regulate the HBV life cycle by impacting on the production of HBV RNAs. This question arose from the observation that 1C8, a compound able to dephosphorylate some RBPs, including SR proteins involved in the HBV life cycle, could down-regulate the production of nascent HBV RNAs [26,47].

Among the kinases identified as the main 1C8 targets *in vitro,* we focused on DYRK1A. This kinase belongs to the DYRK family, within the CMGC group of kinases, a family of constitutively active serine/threonine kinases that participate in a wide variety of processes involved in cell growth, differentiation and transcription [49,50]. DYRK1A is an essential gene in mammals, whose alterations in expression have been associated to different human diseases including Down syndrome when present in trisomy or a neurodevelopmental syndrome when present in haploinsufficiency [65]. It is a pleiotropic kinase regulating a variety of cellular processes including transcription and splicing. DYRK1B is the closest paralog of DYRK1A within the family, sharing a high degree of homology at the primary and secondary structure, with activities connected mostly to cancer development [66]. DYRK1A and DYRK1B have cytoplasmic and nuclear localization but, only DYRK1A localizes to nuclear speckles, the major storage sites of RBPs, and its over-expression induces speckle disassembly [33]. Not surprisingly, DYRK1A can phosphorylate RBPs, among which several SR proteins [52].

Our results show that the genetic manipulation of DYRK1A levels in HBV-infected hepatocytes modulates the production of HBV RNAs without affecting cccDNA levels, suggesting a positive, stimulatory role. This role is less evident for DYRK1B, though our results indicate that the cellular context might be relevant for this kinase. In fact, we found that depletion of DYRK1A induces a compensatory effect on DYRK1B levels, which could partially compensate for DYRK1A loss, thereby affecting the final impact on HBV RNA production. The kinase activity of DYRK1A is involved in the process based on the concurring results of the inhibitory role of small molecule inhibitors and the lack of activity of a kinase-dead version of DYRK1A on HBV parameters.

Previous studies have shown that DYRK1A is recruited to the proximal promoter of several cellular genes characterized by the presence of a common 10 nucleotides motif [29]. We found that DYRK1A associates to the viral genome, cccDNA, in infected hepatocytes. Recruitment to cccDNA could be dependent on the presence of a DYRK1A motif within the enhancer 1/X promoter as shown by the results with reporter assays mutated for the consensus sequence. This potential binding site is adjacent to a previously described interferon-stimulated response element (ISRE) [53], opening the possibility of a functional cross-talk between the activity of DYRK1A and the interferon pathway. The alignment of more than ten thousand HBV sequences derived from all genotypes (https://hbvdb.lyon.inserm.fr/HBVdb/) indicates that this site is conserved among all genotypes, and further support its role as a general regulatory element. These findings do no exclude that DYRK1A may associate to cccDNA via other target sequences. Additional techniques to precisely identify the DYRK1A association site within the cccDNA, such as the recently developed Cut&Run method [67], could confirm our finding. Additionally, targeted mutations in cccDNA should also be performed to determine whether the effect of DYRK1A on HBV life cycle is dependent on its association to the viral genome.

The results showed a good correlation between the activity of DYRK1A and the production of nascent HBV RNAs or the stimulation of reporters containing the DYRK1A motif, supporting a role for the enzymatic activity of the kinase. However, we detected a small stimulatory activity independent of the kinase activity, which has been also observed previously [29]. This could be due to the capacity of DYRK1A to function as a scaffold protein to recruit other factors involved in the processes.

In any case, the Run-On experiments indicated that DYRK1A stimulates HBV transcription. We envision several scenarios that are not mutually exclusive. First, DYRK1A may regulate the phosphorylation status of the RNA Pol II CTD on cccDNA, thereby contributing to increase transcription initiation and/or elongation. Second, transcriptional regulation by DYRK1A could result from its interaction with chromatin associated factors shown to be involved in cccDNA transcriptional regulation, such as the histone acetyl transferase p300/CBP or the transcriptional repressor HP1 [12,27,28,68,69]. Third, DYRK1A may exert its effect on HBV by targeting some cellular RBPs, as it does phosphorylate several splicing-related proteins. Finally, the DYRK1A effect might require some viral factors. In this regard, HBc preferentially associates to CpG island II within cccDNA, a region which partially covers enhancer 1/X promoter, an association that correlates with a permissive epigenetic state [20,21]. Future experiments should investigate whether HBc and DYRK1A can interact in infected hepatocytes as well as consequences of this interaction on HBc phosphorylation and association to viral DNA and RNAs.

A major question raised this study concerns the role of DYRK1A during the HBV life cycle. We are aware that the effect of DYRK1A on the level of HBV RNAs was moderate (always lower than 2-fold), arguing against a major function during the productive viral cycle. However, this level of enhancement may be important for the onset of HBV transcription once the cccDNA is formed in the nucleus, in particular to fire transcription of HBx mRNAs, an event that is required for the initiation of the complete HBV transcriptional program [12,13]. Indeed, since the HBx protein is not present within the viral capsids, some basal level of HBx mRNA synthesis has to occur, early after infection, in order to induce Smc5/6 degradation and de-repress the HBV genome [14–16]. While some studies have reported the presence of HBx mRNA within secreted particles, no evidence has been provided so far to show that these mRNA molecules are transferred into cells and produce HBx [11,70,71]. Therefore, it is possible that a low and transient stimulation of HBx mRNA transcription early upon cccDNA formation may lead to the synthesis of a basal level of HBx sufficient to induce Smc5/6 degradation [72]. In this regard, it is interesting to note that in a recent phospho-proteomic analysis we found that DYRK1A was among the top list of kinases whose activity is predicted to be up-regulated upon HBV infection [73].

In conclusion, this study describes for the first time the involvement of DYRK1A, a kinase which controls the expression of several cellular genes, in the regulation of HBV RNAs production and strongly suggests that this nuclear kinase may be involved in the early control of HBV transcription. Further deciphering the exact role of DYRK1A during the HBV life cycle as well as its molecular targets will be important to further understand the factors which are critical to unleash the HBV transcriptional program and to develop strategies to counteract it.

## ACKNOWLEDGMENTS

We would like to thank Maud Michelet and Anëlle Dubois for their help for PHH isolation, as well as the staff from Pr. Michel Rivoire’s and Dr. Guillaume Passot’s surgery rooms for providing us with liver resections.

## Supplementary

**Supplemental Figure 1.** Kinase profiler assay of 1C8 (10 µM). Only the kinases inhibited by 1C8 with a residual activity<20% are shown. Grey bars indicate the CLK and DYRK kinases targeted by the compound.

**Supplemental Figure 2. RNA levels of selected CMGC kinases in dHepaRG, PHHs and liver biopsies.** RNA levels of CMGC kinases indicated measured by RT-qPCR. Relative RNA levels are expressed by the 2ΔCt value calculated using the PRNP mRNA as control in HepaRG cells (undifferentiated or differentiated, **A**) or PHHs (**B**), HBV-infected or mock-infected. PHHs used were purified from two different donors, each analyzed in triplicate. **C.** CLK1, CLK2, DYRK1A, DYRK1B and SRPK1 mRNA expression in human liver using RNA-Seq data retrieved from Yoo *et al.* [74] expressed as Reads Per Kilobase Million (RPKM).

**Supplemental Figure 3. Comparative effects of AZ191 and DB18 in HBV-infected PHH. A.** Experimental outline. B. PHH were infected and total viral RNAs and pgRNA were quantified by RTqPCR after treatment. IFN: Interferon-α (500 IU/mL). Doses of AZ191 and DB18 were 10 µM and 20 µM. Results are expressed as the mean +/- SD of two independent experiments, each performed in triplicate.

**Supplemental Figure 4. Toxicity assays of AZ191 in PHHs.** Cells infected and treated with various concentrations of AZ191, as indicated in Figure 2A, were analyzed for signs of toxicity using either a neutral red (**A**) or a sulforhodamine B (**B**) assay. Puromycin (Puro) was used as a positive control at 10 µM. Results are expressed as the mean +/- SD of 3 independent experiments, each performed in triplicate.

**Supplemental Figure 5. Effect of CLK2 and SRPK1 KD in HBV-infected dHepaRG cells. A.** Experimental outline. **B.** KD efficiency of the siRNA treatments measured by RT-qPCR at 15-dpt. Results are shown relative to siCTL. **C.** Effect of the kinases KD on the production HBV intracellular RNAs and secreted HBs/HBe antigens, expressed as relative to siCTL (mean +/- SD, n>5, each performed in triplicate).

**Supplemental Figure 6. Toxicity of DYRK1A and DYRK1B KD in HBV-infected dHepaRG cells. A.** Experimental outline. HBV-infected cells were transfected once with siRNA and then harvested at 7-, and 10-dpt. **B** and **C**. Toxicity assays in siRNA transfected dHepaRG cells. Infected and transfected cells were analyzed for signs of toxicity using either a neutral red (**B**) or a sulforhodamine B (**C**) assay. Puromycin (Puro) was used as positive control at 10 µM. In all cases, results are shown relative to siCTL (mean +/- SD, n=2).

**Supplemental Figure 7. Time course analysis of DYRK1A and DYRK1B KD in HBV-infected dHepaRG cells. A.** Experimental outline. HBV-infected cells were transfected once with siRNA and then harvested at 1-, 2-, and 4-dpt. **B** and **C**. Toxicity assays in siRNA transfected dHepaRG cells. Infected and transfected cells were analyzed for signs of toxicity using either a neutral red (**B**) or a sulforhodamine B (**C**) assay. Puromycin (Puro) was used as positive control at 10 µM. **D.** KD efficiency of the siRNA treatments measured by RT-qPCR at the indicated time points. Results are shown relative to siCTL (mean +/- SD, n=2-3). **E**. Effect of the KD for the kinases indicated on HBV RNA production. Intracellular RNA extracted at each time point was analyzed by RT-qPCR to quantify HBV total RNAs. In all cases, results are shown relative to siCTL (mean +/- SD, n=2).

**Supplemental Figure 8. Divergent effects of DYRK1A KD in HBV-infected PHH. A.** Experimental outline. PHH were infected with HBV and 7 days later transfected with siRNA targeting DYRK1A or with a control siRNA (siCTL). **B**. Representative Western blot analysis of DYRK1A KD levels. **C** to **D.** DYRK1A and HBV RNA analysis of a set of 3 independent experiments, each performed in triplicate, in which a decrease of HBV RNAs was observed. **E** and **F** represent another set of 4 experiments in which HBV RNAs increased upon DYRK1A KD. Results are expressed as the mean +/- SD. Statistical analysis is from comparisons between siDYRK1A and control siRNA for each of the measurements.

**Supplemental Figure 9. Compensatory effects of DYRK1A or DYRK1B levels in HBV-infected PHHs and dHepaRG cells**. **A.** mRNA levels were quantified in HBV-infected PHHs transfected with siRNA targeting DYRK1A as shown in Figure S7A. Results were clustered according to their effect on HBV RNA levels (see Figure S7D and F). **B**. Analysis of DYRK1A/DYRK1B compensatory mRNAs variations in HBV-infected dHepaRG cells transfected with siRNA targeting either of the kinases. Statistical analysis is from comparisons between siRNA targeting one of the kinases and control siRNA for DYRK1A or DYRK1B mRNA levels.

**Supplemental Figure 10. Effect of single or combined DYRK1A/DYRK1B KD in HBV-infected PHHs. A.** DYRK1A and DYRK1B protein levels in HBV-infected PHHs with KD for each kinase. Cells were infected with HBV and then transfected with indicated siRNA. Western blot analysis was performed at 7-dpt. Cyclin D1 levels were analyzed as a marker of functional DYRK1 KD, since its accumulation has been shown to depend on both DYRK1A and DYRK1B [57,75]. PHHs (+) and (-) refers to experiments in which an increase (+) or a decrease (-) of HBV RNAs levels was observed following DYRK1A KD (see Figure S7). The bands were quantified using ImageJ (numbers relative to control siRNA, below the blots). **B.** and **C.** Effect of single DYRK1B or double DYRK1A/1B KD in HBV-infected PHHs. Cells were infected and transfected as indicated in Figure S7A. DYRK1A, DYRK1B (**B**), and HBV (**C**) RNAs were quantified at 7-dpt. Results are normalized versus control siRNA and expressed as the mean +/- SD of 2 independent experiments, each performed in triplicate.

**Supplemental Figure 11. Effect of DYRK1B and DYRK1B_KR over-expression on HBV parameters.** dHepaRG cells engineered to express either HA-DYRK1B or HA-DYRK1B_KR under the control of a tetracyclin (Tet)-inducible promoter were infected with HBV and then treated with Tet as indicated in Figure 4A. **A.** Western blot analysis to detect over-expressed HA-DYRK1B or HA-DYRK1B_KR. **B** and **C.** Quantification of HBV RNAs. Results are normalized to the mock situation and expressed as the mean +/- SD, of 2 independent experiments, each performed in triplicate.

## REFERENCES

1. Lucifora J, Protzer U. Attacking hepatitis B virus cccDNA--The holy grail to hepatitis B cure. J Hepatol. 2016;64(1 Suppl):S41–8. doi: 10.1016/j.jhep.2016.02.009. PubMed PMID: 27084036.

2. Zeisel MB, Lucifora J, Mason WS, Sureau C, Beck J, Levrero M, et al. Towards an HBV cure: state-of-the-art and unresolved questions--report of the ANRS workshop on HBV cure. Gut. 2015;64(8):1314–26. doi: 10.1136/gutjnl-2014-308943. PubMed PMID: 25670809.

3. Fanning GC, Zoulim F, Hou J, Bertoletti A. Therapeutic strategies for hepatitis B virus infection: towards a cure. Nat Rev Drug Discov. 2019. Epub 2019/08/29. doi: 10.1038/s41573-019-0037-0. PubMed PMID: 31455905.

4. Martinez MG, Smekalova E, Combe E, Gregoire F, Zoulim F, Testoni B. Gene Editing Technologies to Target HBV cccDNA. Viruses. 2022;14(12). Epub 2022/12/24. doi: 10.3390/v14122654. PubMed PMID: 36560658; PubMed Central PMCID: PMCPMC9787400.

5. Tsukuda S, Watashi K. Hepatitis B virus biology and life cycle. Antiviral Res. 2020;182:104925. Epub 2020/09/01. doi: 10.1016/j.antiviral.2020.104925. PubMed PMID: 32866519.

6. Blondot ML, Bruss V, Kann M. Intracellular transport and egress of hepatitis B virus. J Hepatol. 2016;64(1 Suppl):S49-59. doi: 10.1016/j.jhep.2016.02.008. PubMed PMID: 27084037.

7. Diogo Dias J, Sarica N, Neuveut C. Early Steps of Hepatitis B Life Cycle: From Capsid Nuclear Import to cccDNA Formation. Viruses. 2021;13(5). Epub 2021/05/01. doi: 10.3390/v13050757. PubMed PMID: 33925977; PubMed Central PMCID: PMCPMC8145197.

8. Quasdorff M, Protzer U. Control of hepatitis B virus at the level of transcription. J Viral Hepat. 2010;17(8):527–36. Epub 2010/06/16. doi: 10.1111/j.1365-2893.2010.01315.x. PubMed PMID: 20546497.

9. Xia Y, Guo H. Hepatitis B virus cccDNA: Formation, regulation and therapeutic potential. Antiviral Res. 2020;180:104824. Epub 2020/05/26. doi: 10.1016/j.antiviral.2020.104824. PubMed PMID: 32450266; PubMed Central PMCID: PMCPMC7387223.

10. Sommer G, Heise T. Posttranscriptional control of HBV gene expression. Front Biosci. 2008;13:5533–47. Epub 2008/05/30. doi: 10.2741/3097. PubMed PMID: 18508603.

11. Stadelmayer B, Diederichs A, Chapus F, Rivoire M, Neveu G, Alam A, et al. Full-length 5’RACE identifies all major HBV transcripts in HBV-infected hepatocytes and patient serum. J Hepatol. 2020;73(1):40–51. Epub 2020/02/23. doi: 10.1016/j.jhep.2020.01.028. PubMed PMID: 32087349.

12. Belloni L, Pollicino T, De Nicola F, Guerrieri F, Raffa G, Fanciulli M, et al. Nuclear HBx binds the HBV minichromosome and modifies the epigenetic regulation of cccDNA function. Proc Natl Acad Sci U S A. 2009;106(47):19975–9. doi: 10.1073/pnas.0908365106. PubMed PMID: 19906987; PubMed Central PMCID: PMCPMC2775998.

13. Lucifora J, Arzberger S, Durantel D, Belloni L, Strubin M, Levrero M, et al. Hepatitis B virus X protein is essential to initiate and maintain virus replication after infection. J Hepatol. 2011;55(5):996–1003. Epub 2011/03/08. doi: 10.1016/j.jhep.2011.02.015. PubMed PMID: 21376091.

14. Abdul F, Diman A, Baechler B, Ramakrishnan D, Kornyeyev D, Beran RK, et al. Smc5/6 silences episomal transcription by a three-step function. Nat Struct Mol Biol. 2022;29(9):922–31. Epub 2022/09/14. doi: 10.1038/s41594-022-00829-0. PubMed PMID: 36097294.

15. Decorsiere A, Mueller H, van Breugel PC, Abdul F, Gerossier L, Beran RK, et al. Hepatitis B virus X protein identifies the Smc5/6 complex as a host restriction factor. Nature. 2016;531(7594):386-9. Epub 2016/03/18. doi: 10.1038/nature17170. PubMed PMID: 26983541.

16. Murphy CM, Xu Y, Li F, Nio K, Reszka-Blanco N, Li X, et al. Hepatitis B Virus X Protein Promotes Degradation of SMC5/6 to Enhance HBV Replication. Cell Rep. 2016;16(11):2846–54. Epub 2016/09/15. doi: 10.1016/j.celrep.2016.08.026. PubMed PMID: 27626656; PubMed Central PMCID: PMCPMC5078993.

17. Diab A, Foca A, Zoulim F, Durantel D, Andrisani O. The diverse functions of the hepatitis B core/capsid protein (HBc) in the viral life cycle: Implications for the development of HBc-targeting antivirals. Antiviral Res. 2018;149:211–20. doi: 10.1016/j.antiviral.2017.11.015. PubMed PMID: 29183719; PubMed Central PMCID: PMCPMC5757518.

18. Bock CT, Schranz P, Schroder CH, Zentgraf H. Hepatitis B virus genome is organized into nucleosomes in the nucleus of the infected cell. Virus Genes. 1994;8(3):215–29. Epub 1994/07/01. doi: 10.1007/BF01703079. PubMed PMID: 7975268.

19. Bock CT, Schwinn S, Locarnini S, Fyfe J, Manns MP, Trautwein C, et al. Structural organization of the hepatitis B virus minichromosome. J Mol Biol. 2001;307(1):183–96. Epub 2001/03/13. doi: 10.1006/jmbi.2000.4481. PubMed PMID: 11243813.

20. Chong CK, Cheng CYS, Tsoi SYJ, Huang FY, Liu F, Seto WK, et al. Role of hepatitis B core protein in HBV transcription and recruitment of histone acetyltransferases to cccDNA minichromosome. Antiviral Res. 2017;144:1–7. Epub 2017/05/14. doi: 10.1016/j.antiviral.2017.05.003. PubMed PMID: 28499864.

21. Guo YH, Li YN, Zhao JR, Zhang J, Yan Z. HBc binds to the CpG islands of HBV cccDNA and promotes an epigenetic permissive state. Epigenetics. 2011;6(6):720–6. Epub 2011/05/07. doi: 10.4161/epi.6.6.15815. PubMed PMID: 21546797.

22. Tu T, Zehnder B, Qu B, Urban S. D e novo synthesis of hepatitis B virus nucleocapsids is dispensable for the maintenance and transcriptional regulation of cccDNA. JHEP Rep. 2021;3(1):100195. Epub 2021/01/02. doi: 10.1016/j.jhepr.2020.100195. PubMed PMID: 33385130; PubMed Central PMCID: PMCPMC7771110.

23. Zhong Y, Wu C, Xu Z, Teng Y, Zhao L, Zhao K, et al. Hepatitis B Virus Core Protein Is Not Required for Covalently Closed Circular DNA Transcriptional Regulation. J Virol. 2022;96(21):e0136222. Epub 2022/10/14. doi: 10.1128/jvi.01362-22. PubMed PMID: 36226986; PubMed Central PMCID: PMCPMC9645219.

24. Lucifora J, Pastor F, Charles E, Pons C, Auclair H, Fusil F, et al. Evidence for long-term association of virion-delivered HBV core protein with cccDNA independently of viral protein production. JHEP Rep. 2021;3(5):100330. Epub 2021/08/20. doi: 10.1016/j.jhepr.2021.100330. PubMed PMID: 34409278; PubMed Central PMCID: PMCPMC8363821.

25. Locatelli M, Quivy JP, Chapus F, Michelet M, Fresquet J, Maadadi S, et al. HIRA Supports Hepatitis B Virus Minichromosome Establishment and Transcriptional Activity in Infected Hepatocytes. Cell Mol Gastroenterol Hepatol. 2022;14(3):527–51. Epub 2022/06/02. doi: 10.1016/j.jcmgh.2022.05.007. PubMed PMID: 35643233; PubMed Central PMCID: PMCPMC9304598.

26. Chabrolles H, Auclair H, Vegna S, Lahlali T, Pons C, Michelet M, et al. Hepatitis B virus Core protein nuclear interactome identifies SRSF10 as a host RNA-binding protein restricting HBV RNA production. PLoS Pathog. 2020;16(11):e1008593. Epub 2020/11/13. doi: 10.1371/journal.ppat.1008593. PubMed PMID: 33180834; PubMed Central PMCID: PMCPMC7707522 following competing interests: Uri Lopatin is an advisor to and shareholder of Assembly Biosciences. Lee Arnold was an employee of Assembly Biosciences.

27. Li S, Xu C, Fu Y, Lei PJ, Yao Y, Yang W, et al. DYRK1A interacts with histone acetyl transferase p300 and CBP and localizes to enhancers. Nucleic Acids Res. 2018;46(21):11202–13. Epub 2018/08/24. doi: 10.1093/nar/gky754. PubMed PMID: 30137413; PubMed Central PMCID: PMCPMC6265467.

28. Jang SM, Azebi S, Soubigou G, Muchardt C. DYRK1A phoshorylates histone H3 to differentially regulate the binding of HP1 isoforms and antagonize HP1-mediated transcriptional repression. EMBO Rep. 2014;15(6):686–94. Epub 2014/05/14. doi: 10.15252/embr.201338356. PubMed PMID: 24820035; PubMed Central PMCID: PMCPMC4197879.

29. Di Vona C, Bezdan D, Islam AB, Salichs E, Lopez-Bigas N, Ossowski S, et al. Chromatin-wide profiling of DYRK1A reveals a role as a gene-specific RNA polymerase II CTD kinase. Mol Cell. 2015;57(3):506–20. Epub 2015/01/27. doi: 10.1016/j.molcel.2014.12.026. PubMed PMID: 25620562.

30. Lu H, Yu D, Hansen AS, Ganguly S, Liu R, Heckert A, et al. Phase-separation mechanism for C-terminal hyperphosphorylation of RNA polymerase II. Nature. 2018;558(7709):318-23. Epub 2018/06/01. doi: 10.1038/s41586-018-0174-3. PubMed PMID: 29849146; PubMed Central PMCID: PMCPMC6475116.

31. Gripon P, Rumin S, Urban S, Le Seyec J, Glaise D, Cannie I, et al. Infection of a human hepatoma cell line by hepatitis B virus. Proc Natl Acad Sci U S A. 2002;99(24):15655–60. Epub 2002/11/15. doi: 10.1073/pnas.232137699. PubMed PMID: 12432097; PubMed Central PMCID: PMCPMC137772.

32. Lecluyse EL, Alexandre E. Isolation and culture of primary hepatocytes from resected human liver tissue. Methods Mol Biol. 2010;640:57–82. Epub 2010/07/21. doi: 10.1007/978-1-60761-688-7_3. PubMed PMID: 20645046.

33. Alvarez M, Estivill X, de la Luna S. DYRK1A accumulates in splicing speckles through a novel targeting signal and induces speckle disassembly. J Cell Sci. 2003;116(Pt 15):3099–107. Epub 2003/06/12. doi: 10.1242/jcs.00618. PubMed PMID: 12799418.

34. Michelet M, Alfaiate D, Chardes B, Pons C, Faure-Dupuy S, Engleitner T, et al. Inducers of the NF-kappaB pathways impair hepatitis delta virus replication and strongly decrease progeny infectivity in vitro. JHEP Rep. 2022;4(3):100415. Epub 2022/02/11. doi: 10.1016/j.jhepr.2021.100415. PubMed PMID: 35141510; PubMed Central PMCID: PMCPMC8792426.

35. Ladner SK, Otto MJ, Barker CS, Zaifert K, Wang GH, Guo JT, et al. Inducible expression of human hepatitis B virus (HBV) in stably transfected hepatoblastoma cells: a novel system for screening potential inhibitors of HBV replication. Antimicrob Agents Chemother. 1997;41(8):1715–20. Epub 1997/08/01. PubMed PMID: 9257747; PubMed Central PMCID: PMCPMC163991.

36. Luangsay S, Gruffaz M, Isorce N, Testoni B, Michelet M, Faure-Dupuy S, et al. Early inhibition of hepatocyte innate responses by hepatitis B virus. J Hepatol. 2015;63(6):1314–22. doi: 10.1016/j.jhep.2015.07.014. PubMed PMID: 26216533.

37. Lahlali T, Berke JM, Vergauwen K, Foca A, Vandyck K, Pauwels F, et al. Novel Potent Capsid Assembly Modulators Regulate Multiple Steps of the Hepatitis B Virus Life Cycle. Antimicrob Agents Chemother. 2018;62(10). Epub 2018/07/18. doi: 10.1128/AAC.00835-18. PubMed PMID: 30012770; PubMed Central PMCID: PMCPMC6153789.

38. Mueller H, Wildum S, Luangsay S, Walther J, Lopez A, Tropberger P, et al. A novel orally available small molecule that inhibits hepatitis B virus expression. J Hepatol. 2018;68(3):412–20. Epub 2017/10/29. doi: 10.1016/j.jhep.2017.10.014. PubMed PMID: 29079285.

39. Brahmaiah D, Kanaka Durga Bhavani A, Aparna P, Sampath Kumar N, Solhi H, Le Guevel R, et al. Discovery of DB18, a potent inhibitor of CLK kinases with a high selectivity against DYRK1A kinase. Bioorg Med Chem. 2021;31:115962. Epub 2021/01/11. doi: 10.1016/j.bmc.2020.115962. PubMed PMID: 33422908.

40. Isorce N, Lucifora J, Zoulim F, Durantel D. Immune-modulators to combat hepatitis B virus infection: From IFN-alpha to novel investigational immunotherapeutic strategies. Antiviral Res. 2015;122:69–81. doi: 10.1016/j.antiviral.2015.08.008. PubMed PMID: 26275801.

41. Allweiss L, Testoni B, Yu M, Lucifora J, Ko C, Qu B, et al. Quantification of the hepatitis B virus cccDNA: evidence-based guidelines for monitoring the key obstacle of HBV cure. Gut. 2023. Epub 2023/01/28. doi: 10.1136/gutjnl-2022-328380. PubMed PMID: 36707234.

42. Lucifora J, Xia Y, Reisinger F, Zhang K, Stadler D, Cheng X, et al. Specific and nonhepatotoxic degradation of nuclear hepatitis B virus cccDNA. Science. 2014;343(6176):1221-8. doi: 10.1126/science.1243462. PubMed PMID: 24557838.

43. Werle-Lapostolle B, Bowden S, Locarnini S, Wursthorn K, Petersen J, Lau G, et al. Persistence of cccDNA during the natural history of chronic hepatitis B and decline during adefovir dipivoxil therapy. Gastroenterology. 2004;126(7):1750–8. Epub 2004/06/10. doi: 10.1053/j.gastro.2004.03.018. PubMed PMID: 15188170.

44. Mueller H, Lopez A, Tropberger P, Wildum S, Schmaler J, Pedersen L, et al. PAPD5/7 Are Host Factors That Are Required for Hepatitis B Virus RNA Stabilization. Hepatology. 2019;69(4):1398–411. Epub 2018/10/27. doi: 10.1002/hep.30329. PubMed PMID: 30365161.

45. Paulsen MT, Veloso A, Prasad J, Bedi K, Ljungman EA, Magnuson B, et al. Use of Bru-Seq and BruChase-Seq for genome-wide assessment of the synthesis and stability of RNA. Methods. 2014;67(1):45–54. Epub 2013/08/27. doi: 10.1016/j.ymeth.2013.08.015. PubMed PMID: 23973811; PubMed Central PMCID: PMCPMC4009065.

46. Cheung PK, Horhant D, Bandy LE, Zamiri M, Rabea SM, Karagiosov SK, et al. A Parallel Synthesis Approach to the Identification of Novel Diheteroarylamide-Based Compounds Blocking HIV Replication: Potential Inhibitors of HIV-1 Pre-mRNA Alternative Splicing. J Med Chem. 2016;59(5):1869–79. Epub 2016/02/16. doi: 10.1021/acs.jmedchem.5b01357. PubMed PMID: 26878150.

47. Shkreta L, Blanchette M, Toutant J, Wilhelm E, Bell B, Story BA, et al. Modulation of the splicing regulatory function of SRSF10 by a novel compound that impairs HIV-1 replication. Nucleic Acids Res. 2017;45(7):4051–67. doi: 10.1093/nar/gkw1223. PubMed PMID: 27928057; PubMed Central PMCID: PMCPMC5397194.

48. Shkreta L, Toutant J, Delannoy A, Durantel D, Salvetti A, Ehresmann S, et al. The anticancer potential of the CLK kinases inhibitors 1C8 and GPS167 revealed by their impact on the epithelial-mesenchymal transition and the antiviral immune response. Oncotarget. 2024;15:313–25. Epub 2024/05/16. doi: 10.18632/oncotarget.28585. PubMed PMID: 38753413; PubMed Central PMCID: PMCPMC11098031.

49. Aranda S, Laguna A, de la Luna S. DYRK family of protein kinases: evolutionary relationships, biochemical properties, and functional roles. FASEB J. 2011;25(2):449–62. Epub 2010/11/05. doi: 10.1096/fj.10-165837. PubMed PMID: 21048044.

50. Lindberg MF, Meijer L. Dual-Specificity, Tyrosine Phosphorylation-Regulated Kinases (DYRKs) and cdc2-Like Kinases (CLKs) in Human Disease, an Overview. Int J Mol Sci. 2021;22(11). Epub 2021/07/03. doi: 10.3390/ijms22116047. PubMed PMID: 34205123; PubMed Central PMCID: PMCPMC8199962.

51. Wang E, Pineda JMB, Kim WJ, Chen S, Bourcier J, Stahl M, et al. Modulation of RNA splicing enhances response to BCL2 inhibition in leukemia. Cancer Cell. 2023;41(1):164–80 e8. Epub 2022/12/24. doi: 10.1016/j.ccell.2022.12.002. PubMed PMID: 36563682; PubMed Central PMCID: PMCPMC9839614.

52. Pastor F, Shkreta L, Chabot B, Durantel D, Salvetti A. Interplay Between CMGC Kinases Targeting SR Proteins and Viral Replication: Splicing and Beyond. Front Microbiol. 2021;12:658721. Epub 2021/04/16. doi: 10.3389/fmicb.2021.658721. PubMed PMID: 33854493; PubMed Central PMCID: PMCPMC8040976.

53. Belloni L, Allweiss L, Guerrieri F, Pediconi N, Volz T, Pollicino T, et al. IFN-alpha inhibits HBV transcription and replication in cell culture and in humanized mice by targeting the epigenetic regulation of the nuclear cccDNA minichromosome. J Clin Invest. 2012;122(2):529–37. doi: 10.1172/JCI58847. PubMed PMID: 22251702; PubMed Central PMCID: PMCPMC3266786.

54. Schlicksup CJ, Wang JC, Francis S, Venkatakrishnan B, Turner WW, VanNieuwenhze M, et al. Hepatitis B virus core protein allosteric modulators can distort and disrupt intact capsids. Elife. 2018;7. doi: 10.7554/eLife.31473. PubMed PMID: 29377794; PubMed Central PMCID: PMCPMC5788503.

55. Fukuhara T, Hosoya T, Shimizu S, Sumi K, Oshiro T, Yoshinaka Y, et al. Utilization of host SR protein kinases and RNA-splicing machinery during viral replication. Proc Natl Acad Sci U S A. 2006;103(30):11329–33. Epub 2006/07/15. doi: 10.1073/pnas.0604616103. PubMed PMID: 16840555; PubMed Central PMCID: PMCPMC1544086.

56. Muraki M, Ohkawara B, Hosoya T, Onogi H, Koizumi J, Koizumi T, et al. Manipulation of alternative splicing by a newly developed inhibitor of Clks. J Biol Chem. 2004;279(23):24246–54. Epub 2004/03/11. doi: 10.1074/jbc.M314298200. PubMed PMID: 15010457.

57. Ashford AL, Oxley D, Kettle J, Hudson K, Guichard S, Cook SJ, et al. A novel DYRK1B inhibitor AZ191 demonstrates that DYRK1B acts independently of GSK3beta to phosphorylate cyclin D1 at Thr(286), not Thr(288). Biochem J. 2014;457(1):43–56. Epub 2013/10/19. doi: 10.1042/BJ20130461. PubMed PMID: 24134204.

58. Ninomiya K, Kataoka N, Hagiwara M. Stress-responsive maturation of Clk1/4 pre-mRNAs promotes phosphorylation of SR splicing factor. J Cell Biol. 2011;195(1):27–40. Epub 2011/09/29. doi: 10.1083/jcb.201107093. PubMed PMID: 21949414; PubMed Central PMCID: PMCPMC3187705.

59. Ackeifi C, Swartz E, Kumar K, Liu H, Chalada S, Karakose E, et al. Pharmacologic and genetic approaches define human pancreatic beta cell mitogenic targets of DYRK1A inhibitors. JCI Insight. 2020;5(1). Epub 2019/12/11. doi: 10.1172/jci.insight.132594. PubMed PMID: 31821176; PubMed Central PMCID: PMCPMC7030849.

60. Di Vona C, Barba L, Ferrari R, de la Luna S. Loss of the DYRK1A Protein Kinase Results in the Reduction in Ribosomal Protein Gene Expression, Ribosome Mass and Reduced Translation. Biomolecules. 2023;14(1). Epub 2024/01/23. doi: 10.3390/biom14010031. PubMed PMID: 38254631; PubMed Central PMCID: PMCPMC10813206.

61. Ludgate L, Ning X, Nguyen DH, Adams C, Mentzer L, Hu J. Cyclin-dependent kinase 2 phosphorylates s/t-p sites in the hepadnavirus core protein C-terminal domain and is incorporated into viral capsids. J Virol. 2012;86(22):12237–50. doi: 10.1128/JVI.01218-12. PubMed PMID: 22951823; PubMed Central PMCID: PMCPMC3486511.

62. Luo J, Xi J, Gao L, Hu J. Role of Hepatitis B virus capsid phosphorylation in nucleocapsid disassembly and covalently closed circular DNA formation. PLoS Pathog. 2020;16(3):e1008459. Epub 2020/04/01. doi: 10.1371/journal.ppat.1008459. PubMed PMID: 32226051; PubMed Central PMCID: PMCPMC7145273.

63. Diab A, Foca A, Fusil F, Lahlali T, Jalaguier P, Amirache F, et al. Polo-like-kinase 1 is a proviral host factor for hepatitis B virus replication. Hepatology. 2017;66(6):1750–65. doi: 10.1002/hep.29236. PubMed PMID: 28445592; PubMed Central PMCID: PMCPMC5658273.

64. Wittkop L, Schwarz A, Cassany A, Grun-Bernhard S, Delaleau M, Rabe B, et al. Inhibition of protein kinase C phosphorylation of hepatitis B virus capsids inhibits virion formation and causes intracellular capsid accumulation. Cell Microbiol. 2010;12(7):962–75. Epub 2010/01/30. doi: 10.1111/j.1462-5822.2010.01444.x. PubMed PMID: 20109160.

65. Deboever E, Fistrovich A, Hulme C, Dunckley T. The Omnipresence of DYRK1A in Human Diseases. Int J Mol Sci. 2022;23(16). Epub 2022/08/27. doi: 10.3390/ijms23169355. PubMed PMID: 36012629; PubMed Central PMCID: PMCPMC9408930.

66. Becker W. A wake-up call to quiescent cancer cells - potential use of DYRK1B inhibitors in cancer therapy. FEBS J. 2018;285(7):1203–11. Epub 2017/12/02. doi: 10.1111/febs.14347. PubMed PMID: 29193696.

67. Meers MP, Bryson TD, Henikoff JG, Henikoff S. Improved CUT&RUN chromatin profiling tools. Elife. 2019;8. Epub 2019/06/25. doi: 10.7554/eLife.46314. PubMed PMID: 31232687; PubMed Central PMCID: PMCPMC6598765.

68. Cougot D, Wu Y, Cairo S, Caramel J, Renard CA, Levy L, et al. The hepatitis B virus X protein functionally interacts with CREB-binding protein/p300 in the regulation of CREB-mediated transcription. J Biol Chem. 2007;282(7):4277–87. Epub 2006/12/13. doi: 10.1074/jbc.M606774200. PubMed PMID: 17158882.

69. Riviere L, Gerossier L, Ducroux A, Dion S, Deng Q, Michel ML, et al. HBx relieves chromatin-mediated transcriptional repression of hepatitis B viral cccDNA involving SETDB1 histone methyltransferase. J Hepatol. 2015;63(5):1093–102. doi: 10.1016/j.jhep.2015.06.023. PubMed PMID: 26143443.

70. Bai L, Zhang X, Kozlowski M, Li W, Wu M, Liu J, et al. Extracellular Hepatitis B Virus RNAs Are Heterogeneous in Length and Circulate as Capsid-Antibody Complexes in Addition to Virions in Chronic Hepatitis B Patients. J Virol. 2018;92(24). Epub 2018/10/05. doi: 10.1128/JVI.00798-18. PubMed PMID: 30282709; PubMed Central PMCID: PMCPMC6258948.

71. Niu C, Livingston CM, Li L, Beran RK, Daffis S, Ramakrishnan D, et al. The Smc5/6 Complex Restricts HBV when Localized to ND10 without Inducing an Innate Immune Response and Is Counteracted by the HBV X Protein Shortly after Infection. PLoS One. 2017;12(1):e0169648. Epub 2017/01/18. doi: 10.1371/journal.pone.0169648. PubMed PMID: 28095508; PubMed Central PMCID: PMCPMC5240991 does not alter our adherence to PLOS ONE policies on sharing data and materials.

72. Doitsh G, Shaul Y. Enhancer I predominance in hepatitis B virus gene expression. Mol Cell Biol. 2004;24(4):1799–808. Epub 2004/01/30. doi: 10.1128/MCB.24.4.1799-1808.2004. PubMed PMID: 14749394; PubMed Central PMCID: PMCPMC344184.

73. Pastor F, Charles E, Belmudes L, Chabrolles H, Cescato M, Rivoire M, et al. Deciphering the phospho-signature induced by hepatitis B virus in primary human hepatocytes. Front Microbiol. 2024;15:1415449. Epub 2024/06/06. doi: 10.3389/fmicb.2024.1415449. PubMed PMID: 38841065; PubMed Central PMCID: PMCPMC11150682.

74. Yoo S, Wang W, Wang Q, Fiel MI, Lee E, Hiotis SP, et al. A pilot systematic genomic comparison of recurrence risks of hepatitis B virus-associated hepatocellular carcinoma with low- and high-degree liver fibrosis. BMC Med. 2017;15(1):214. Epub 2017/12/08. doi: 10.1186/s12916-017-0973-7. PubMed PMID: 29212479; PubMed Central PMCID: PMCPMC5719570.

75. Chen JY, Lin JR, Tsai FC, Meyer T. Dosage of Dyrk1a shifts cells within a p21-cyclin D1 signaling map to control the decision to enter the cell cycle. Mol Cell. 2013;52(1):87–100. Epub 2013/10/15. doi: 10.1016/j.molcel.2013.09.009. PubMed PMID: 24119401; PubMed Central PMCID: PMCPMC4039290.

