## Supplementary figures and images for "THE DUAL-SPECIFICITY KINASE DYRK1A INTERACTS WITH THE HEPATITIS B VIRUS GENOME AND REGULATES THE PRODUCTION OF VIRAL RNA"

### Figure S1

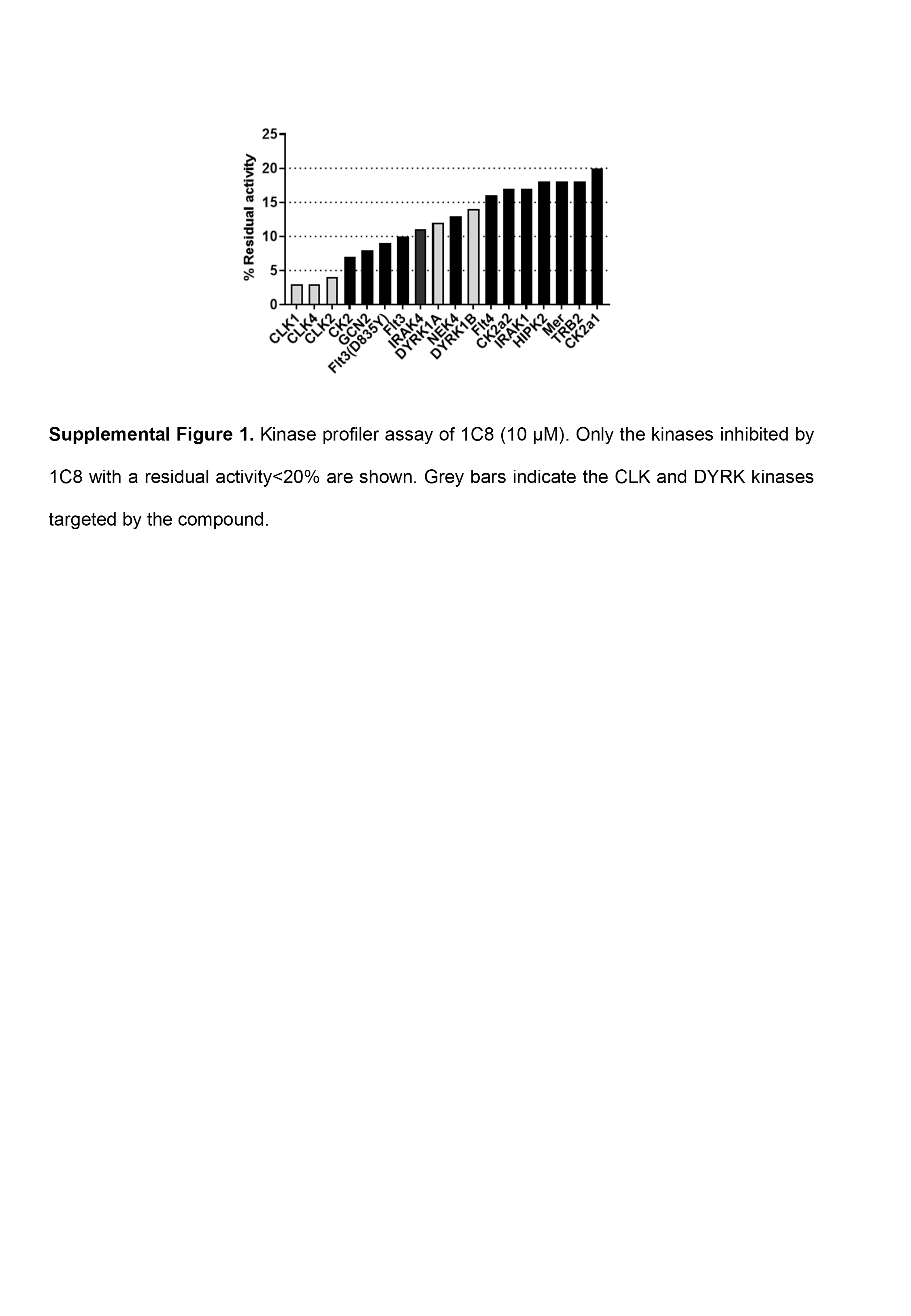

### Figure S2

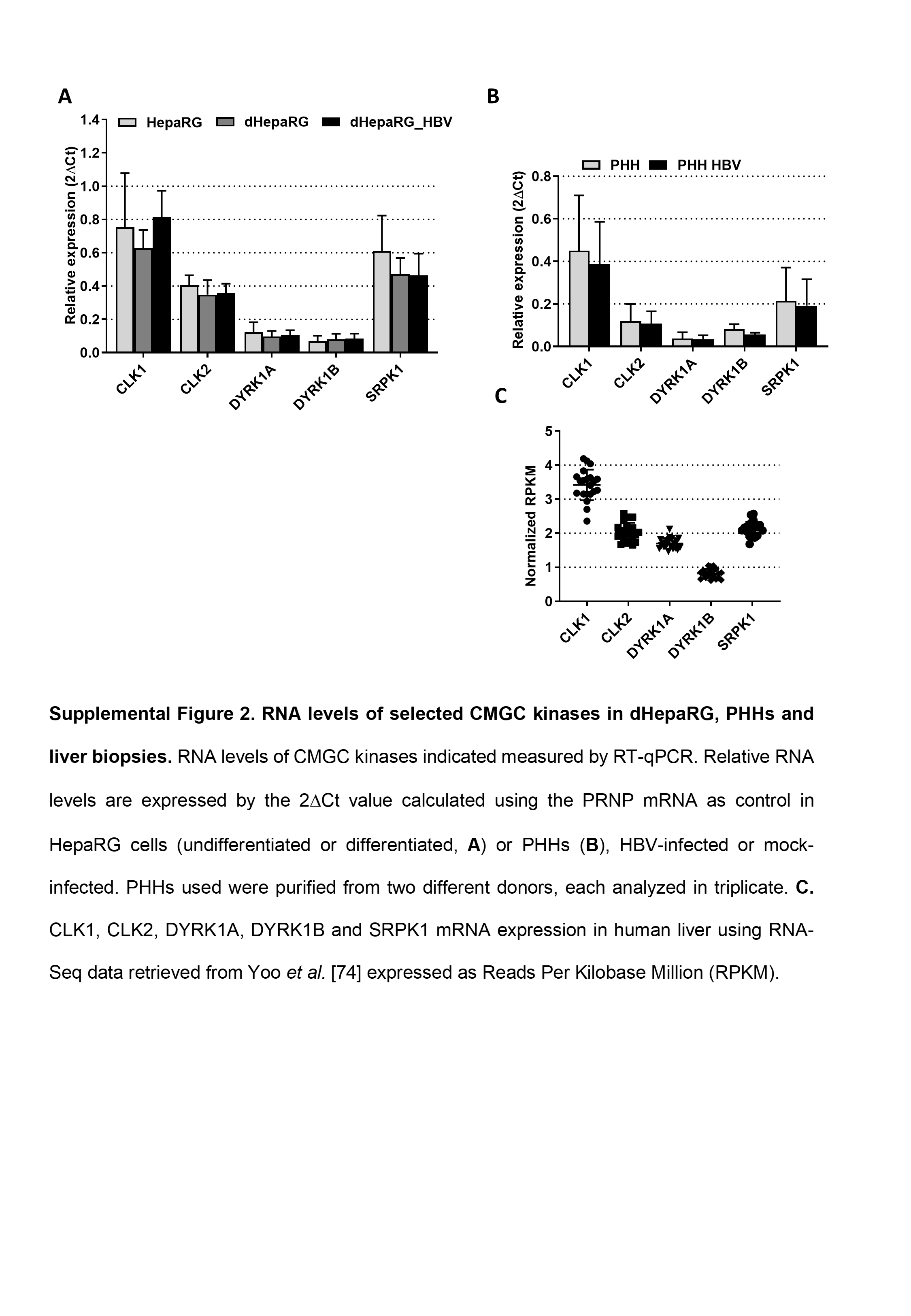

### Figure S3

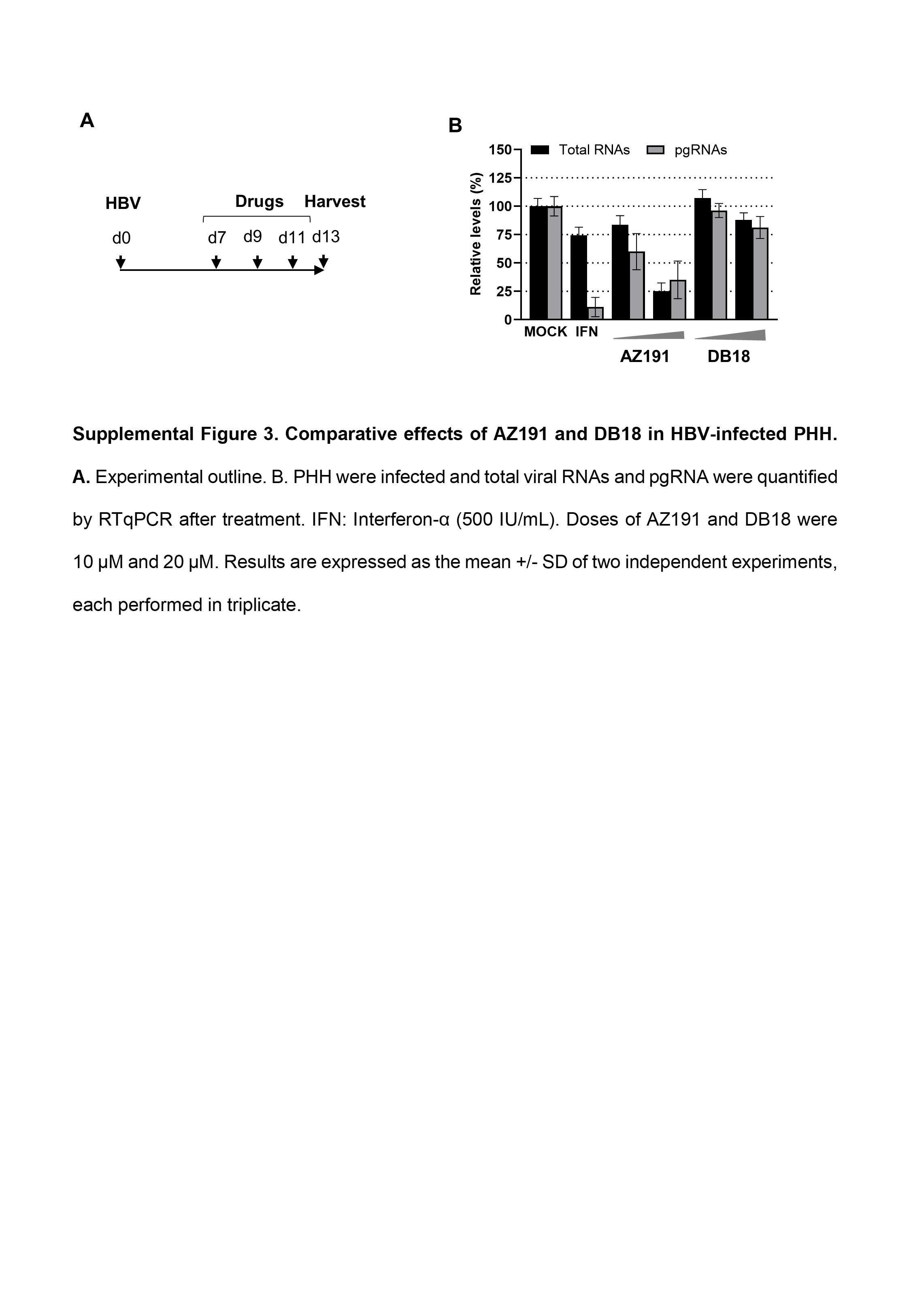

### Figure S4

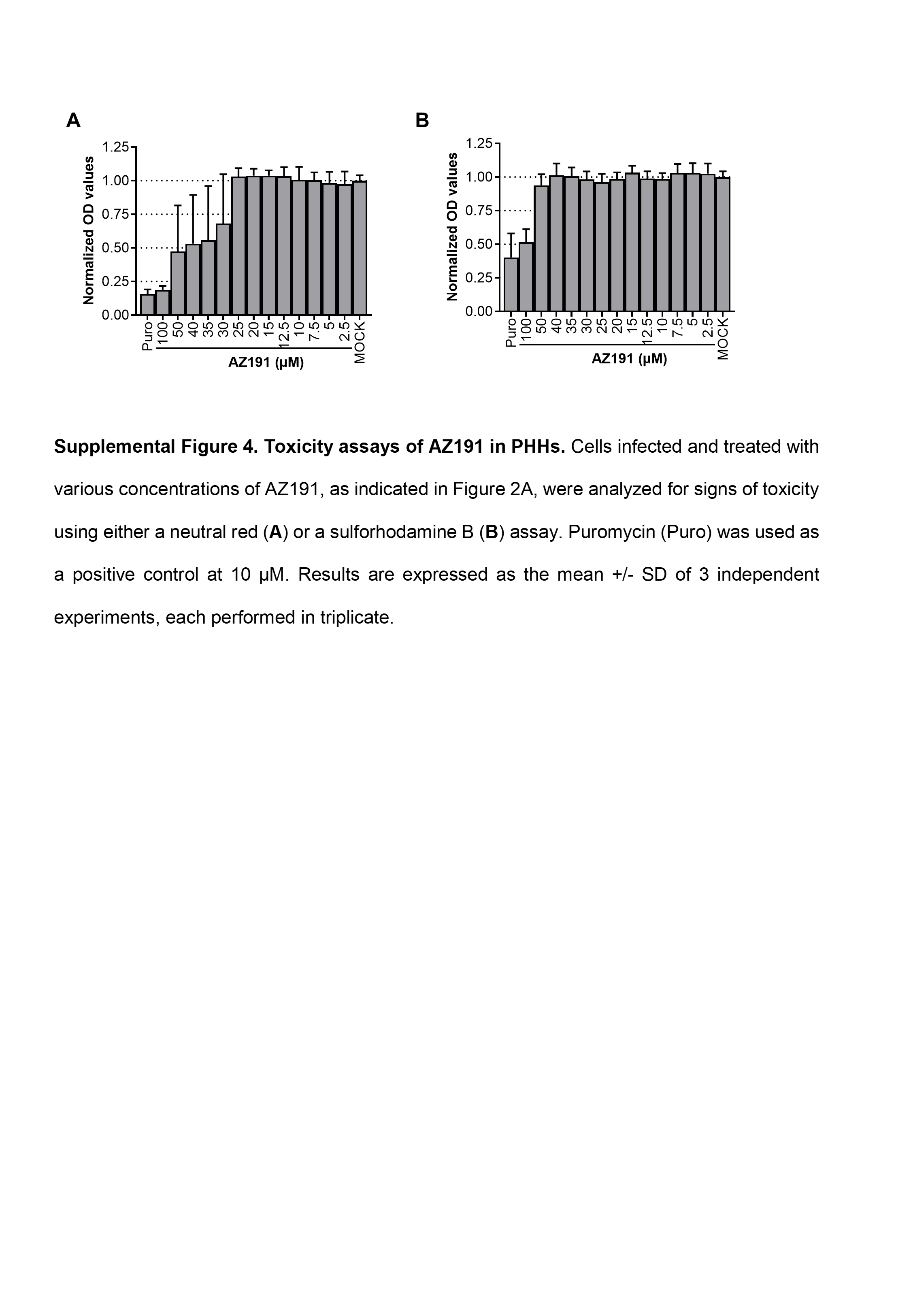

### Figure S5

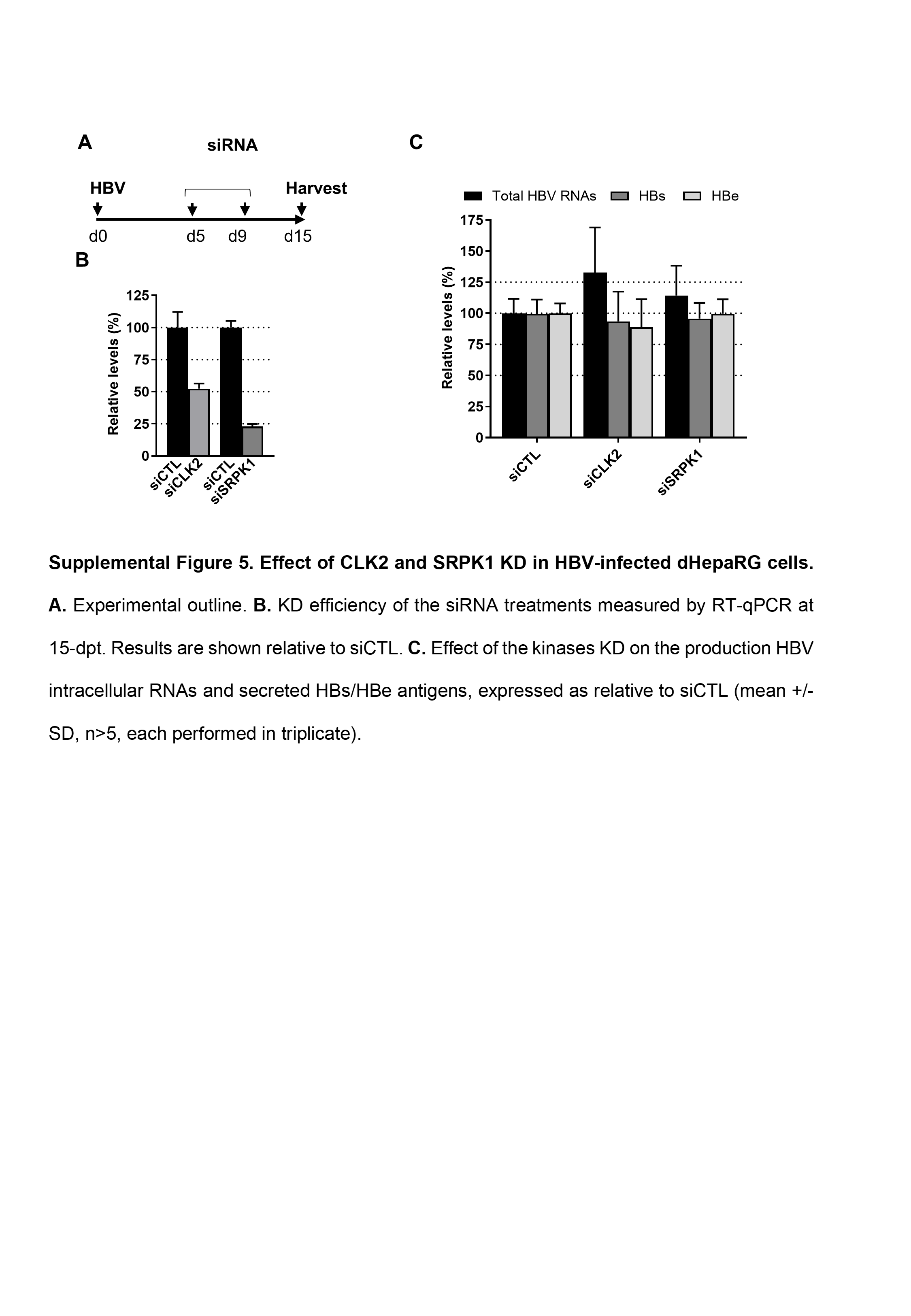

### Figure S6

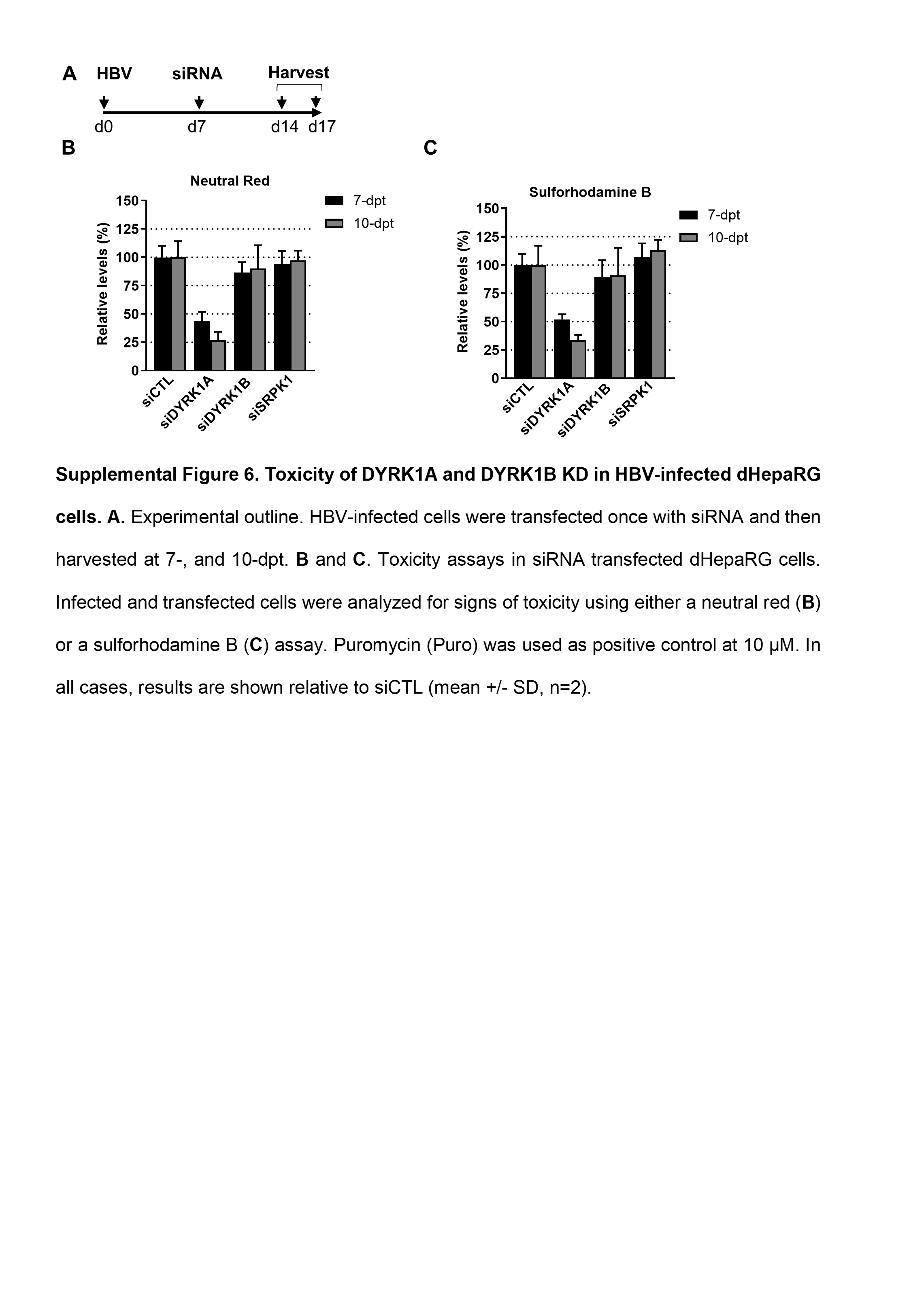

### Figure S7

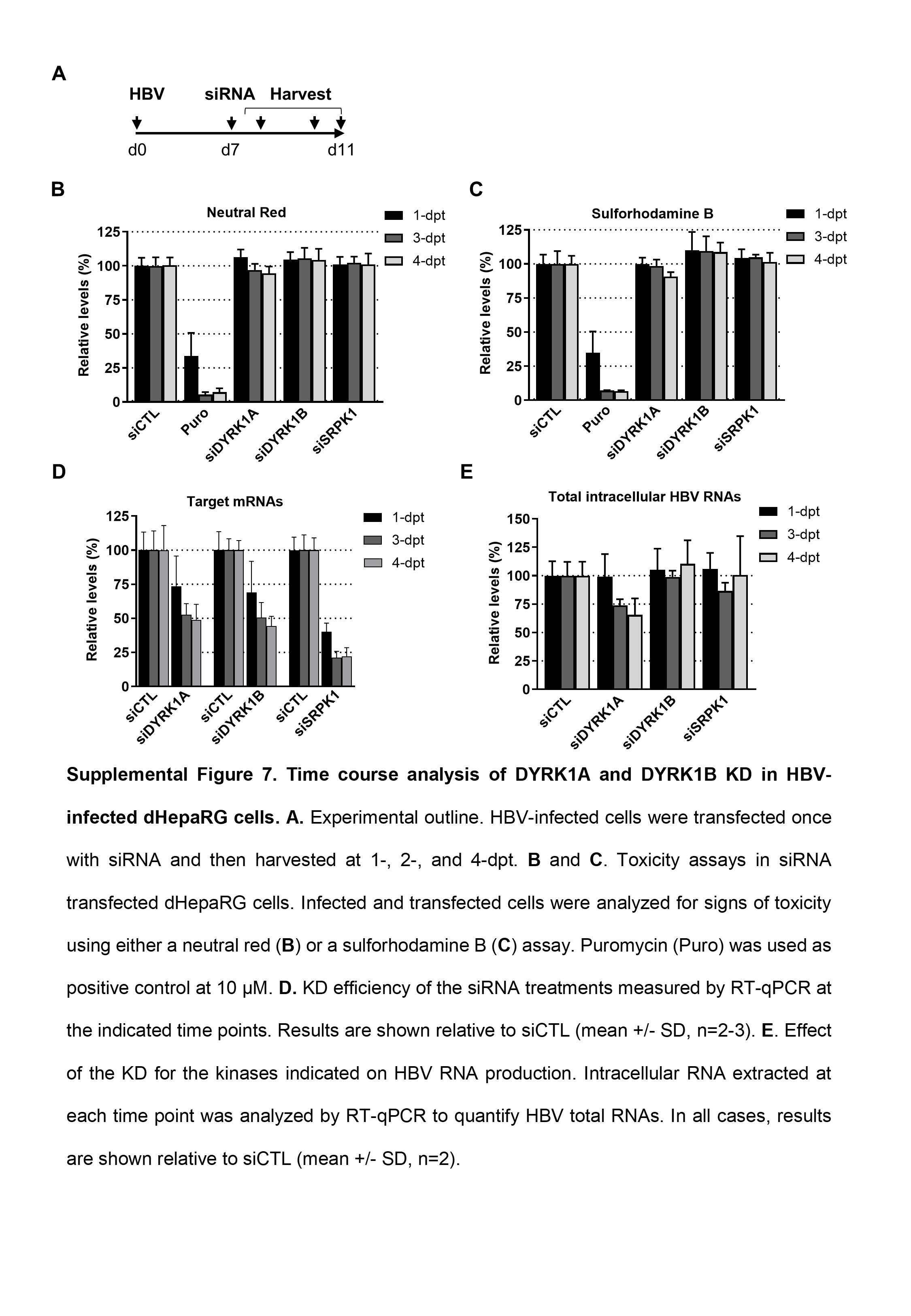

### Figure S8

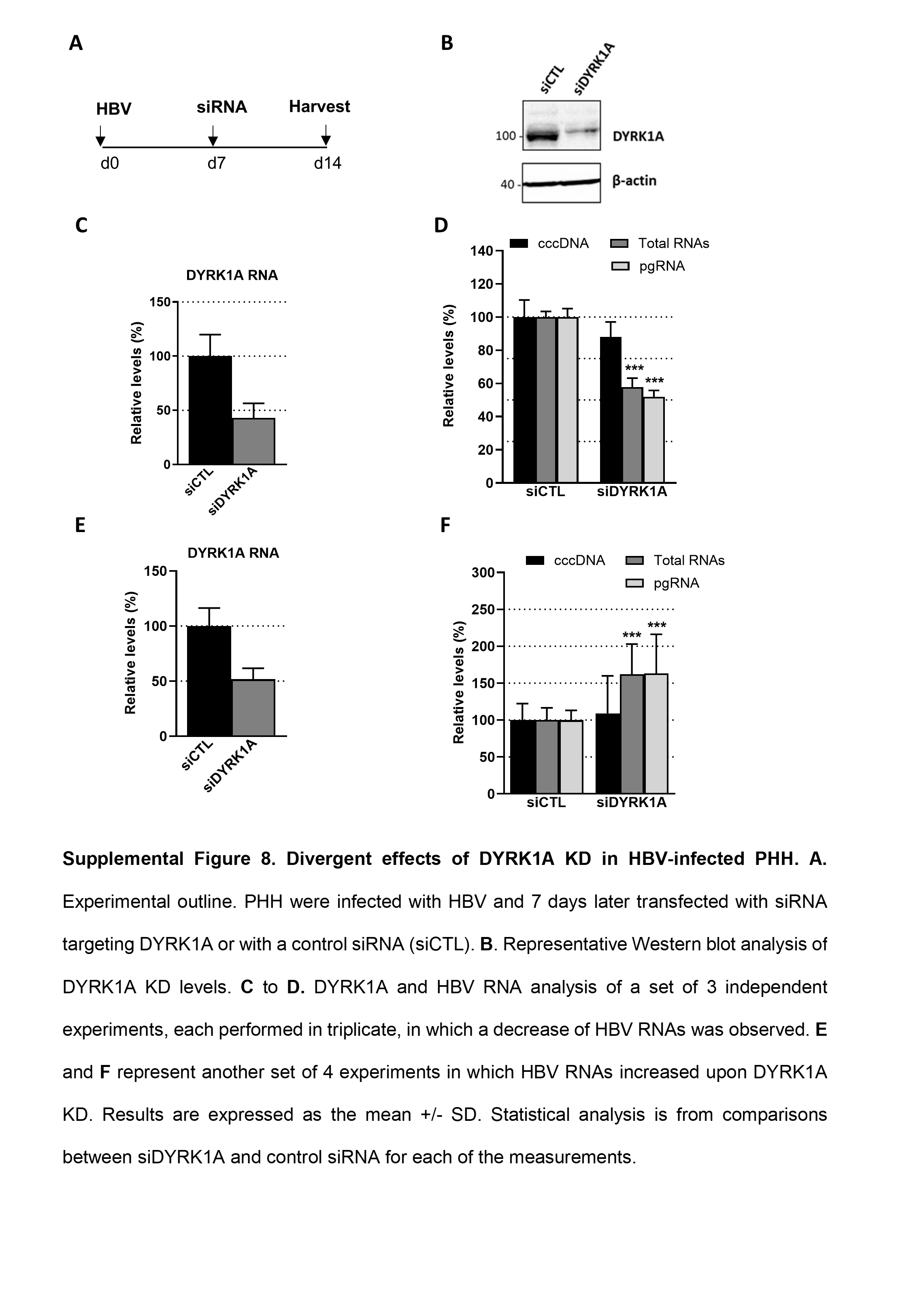

### Figure S9

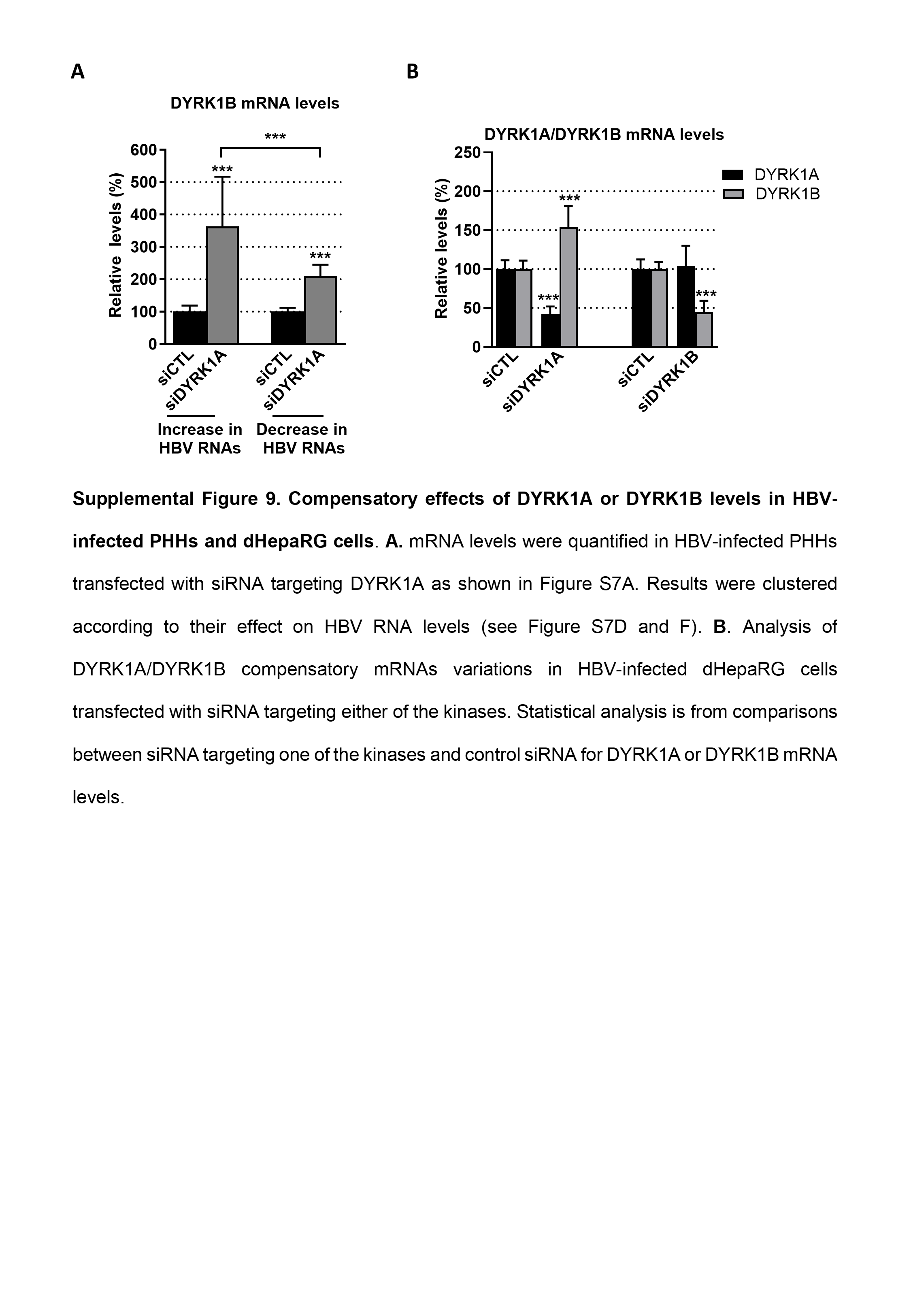

### Figure S10

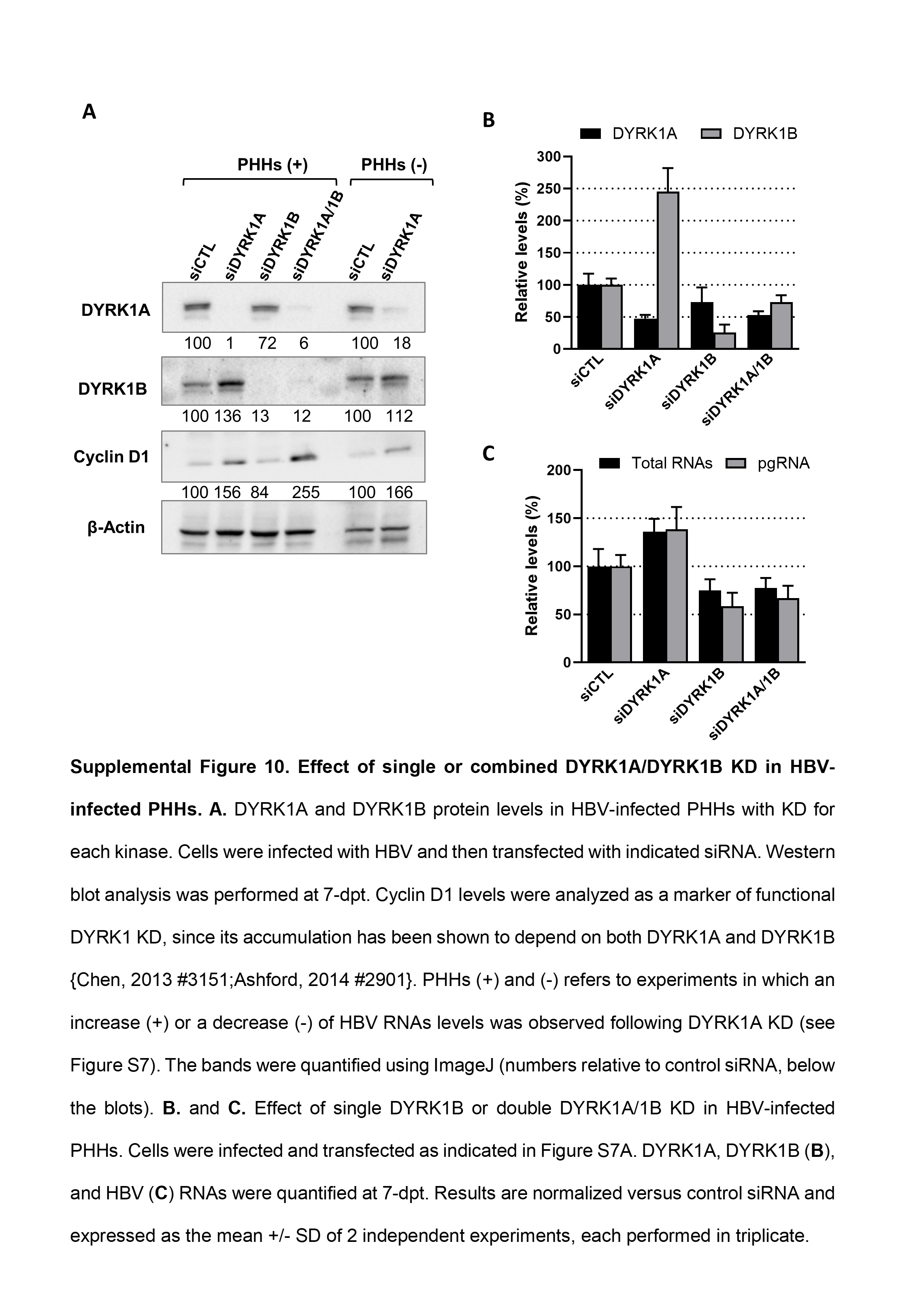

### Figure S11

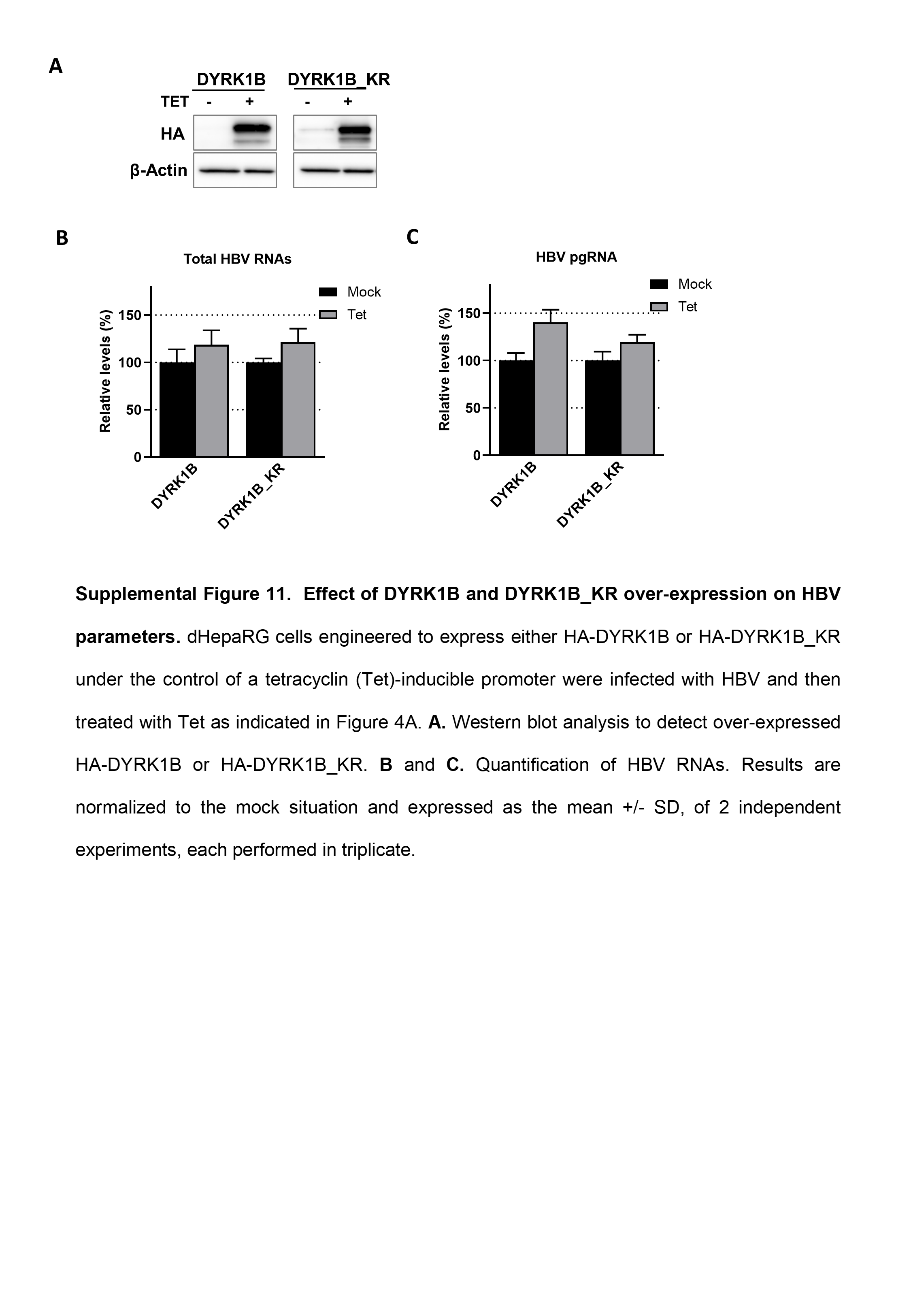
